# Chemogenetic modulation and single-photon calcium imaging in anterior cingulate cortex reveal a mechanism for effort-based decisions

**DOI:** 10.1101/792069

**Authors:** Evan E. Hart, Garrett J. Blair, Thomas J. O’Dell, Hugh T. Blair, Alicia Izquierdo

## Abstract

The anterior cingulate cortex (ACC) is implicated in effort exertion and choices based on effort cost, but it is still unclear how it mediates this cost-benefit evaluation. Here, male rats were trained to exert effort for a high-value reward (sucrose pellets) in a progressive ratio lever pressing task. Trained rats were then tested in two conditions: a no-choice condition where lever pressing for sucrose was the only available food option, and a choice condition where a low-value reward (lab chow) was freely available as an alternative to pressing for sucrose. Disruption of ACC—via either chemogenetic inhibition or excitation—reduced lever pressing in the choice, but not in the no-choice, condition. We next looked for value coding cells in ACC during effortful behavior and reward consumption phases during choice and no-choice conditions. For this, we utilized *in vivo* miniaturized fluorescence microscopy to reliably track responses of the same cells and compare how ACC neurons respond during the same effortful behavior where there was a choice versus when there was no-choice. We found that lever-press and sucrose-evoked responses were significantly weaker during choice compared to no-choice sessions, which may have rendered them more susceptible to chemogenetic disruption. Taken together, findings from our interference experiments and neural recordings suggest that a mechanism by which ACC mediates effortful decisions is in the discrimination of the utility of available options. ACC regulates these choices by providing a stable population code for the relative value of different options.

**Significance Statement:** The anterior cingulate cortex (ACC) is implicated in effort-based decision making. Here, we used chemogenetics and *in vivo* calcium imaging to explore its mechanism. Rats were trained to lever press for a high-value reward and tested in two conditions: a no-choice condition where lever pressing for the high-value reward was the only option, and a choice condition where a low-value reward was also available. Inhibition or excitation of ACC reduced effort toward the high value option, but only in the choice condition. Neural responses in ACC were weaker in the choice compared to the no-choice condition. A mechanism by which ACC regulates effortful decisions is in providing a stable population code for the discrimination of the utility of available options.

## INTRODUCTION

Real-world decisions rarely involve choosing between unambiguously favorable vs. unfavorable options. Often, options must be evaluated along multiple dimensions that incorporate an evaluation of the rewards themselves as well as the actions or efforts to procure them (Skvortsova et al., 2014). For example, we typically make decisions between options in comparison, where one outcome may be more costly (i.e. more effortful) yet more preferred than the other.

Outside of the striatum, which has been the major region of study in such effort-based decision making studies (Cousins et al., 1996; Ghods-Sharifi and Floresco, 2010; Nowend et al., 2001; Salamone et al., 2007; Salamone et al., 2003; Salamone et al., 1994; Salamone et al., 1991), the anterior cingulate cortex (ACC) is also involved in the evaluation of both physical and cognitive effort costs (Cowen et al., 2012; Floresco and Ghods-Sharifi, 2007; Hillman and Bilkey, 2010, 2012; Hosking et al., 2014; Schweimer and Hauber, 2005; Walton et al., 2003; Winstanley and Floresco, 2016). Neurons in both rat and primate ACC signal value during economic decision-making (Azab and Hayden, 2017; Hunt and Hayden, 2017; Lapish et al., 2008; Mashhoori et al., 2018). Neural responses in this region also track trial-by-trial outcomes of choices (Akam et al., 2017; Procyk et al., 2000; Seo and Lee, 2007; Shidara and Richmond, 2002), reward history (Bernacchia et al., 2011), reward prediction errors, (Hayden et al., 2009; Hyman et al., 2017; Kennerley et al., 2011), and counterfactual options (‘rewards not taken’) (Hayden et al., 2009; Mashhoori et al., 2018). Indeed, both anatomical and functional evidence supports the idea that ACC activity supports representations of reward value and action outcomes (Heilbronner and Hayden, 2016; Shenhav et al., 2013).

A paradigm that involves selecting between qualitatively different reinforcers may closely model human decisions where we encounter options that are more/less preferred, not more/less of the same reward identity (Cousins and Salamone, 1994; Nowend et al., 2001; Nunes et al., 2013; Randall et al., 2014a; Randall et al., 2015; Randall et al., 2012; Salamone et al., 2007; Salamone et al., 2017; Salamone et al., 1991; Yohn et al., 2016a; Yohn et al., 2016b; Yohn et al., 2016c). The majority of the seminal rodent studies probing ACC in effort-based choice (Floresco and Ghods-Sharifi, 2007; Hauber and Sommer, 2009; Walton et al., 2003; Walton et al., 2002; Winstanley and Floresco, 2016) have used traditional pharmacological, lesion, and electrophysiological approaches, so a fine-grained analysis involving cell-type specific, temporally-restricted targeting of ACC in effort choice has not yet been reported.

Here, we tested the effects of inhibitory (hM4Di, or G_i_) and excitatory (hM3Dq, or G_q_) Designer Receptors Exclusively Activated by Designer Drugs (DREADDs) (Alexander et al., 2009; Armbruster et al., 2007; Roth, 2016) in ACC on the same effortful choice task that we previously probed following lesions (Hart et al., 2017) and pharmacological inactivations (Hart and Izquierdo, 2017). Briefly, our task required rats to choose between working for a preferred reward (sucrose) vs. consuming a concurrently and freely-available, but less preferred reward (standard chow). We assessed the role of ACC on (i) progressive ratio (PR) lever pressing for sucrose pellets (i.e. general motivation), and (ii) PR lever pressing with choice (PRC) of a freely-available alternative (i.e. effortful decision-making: choosing between working for sucrose pellets vs. concurrently available laboratory chow). We also tested the effects of ACC inhibition and excitation on the choice between sucrose pellets vs. chow when these reinforcers were both freely available (i.e. “free choice”). Finally, in a separate cohort of animals we looked for value coding cells in ACC during effortful behavior and reward consumption phases. For this, we utilized *in vivo* miniaturized fluorescence microscopy (UCLA “miniscopes”, (Ghosh et al., 2011; Aharoni et al., 2019)) to reliably track responses of the same cells and compare how ACC neurons respond during the same effortful behavior in separate sessions where there was a choice (PRC) versus when there was no choice (PR).

## MATERIALS AND METHODS

### Subjects

Subjects were N=44 adult male Long-Evans rats (n=12 G_i_ DREADD experiment, n=12 G_q_ DREADD experiment, n=10 GFP (null virus) control experiment, n=4 calcium imaging experiment, n=6 acute slice recording for validation of DREADDs). Rats were obtained from Charles River Laboratories (Hollister, CA), were PND 60 at the time of arrival to the UCLA vivarium, and were singly-housed for all phases of experiments with the exception of the acclimation period and handling, during which they were pair-housed. All rats were handled for 10 minutes in pairs for 5 d after a brief acclimation period (3 d). Rats weighed an average of 309.1 g at the beginning of experiments. Three subjects were not included in the final data analyses: one from G_i_ experiment and one from G_q_ experiment due to unilateral (not bilateral) viral expression, and one from the GCaMP experiments due to poor stability of imaging over days.

The vivarium was maintained under a 12/12 h reverse light cycle at 22°C, and lab chow and water were available *ad libitum* prior to behavioral testing. Rats were food restricted one day prior to behavioral testing to ensure motivation to work for rewards. Given the sensitivity of the behavioral tests on motivation, special care was taken to maintain consistent food rations throughout the experiment. This was 12 g/d at the beginning of testing, but then decreased to 8 g/d at the beginning of the choice phase (details below). Rats were monitored every other day for their body weight, and were never permitted to drop below their 85% free feeding baseline weight. Training and testing were conducted during the early portion of the dark cycle (∼0800 to 1200 H). Experiments were conducted 5-7 d per week, and rats were fed once daily on weekends (12 g) when testing was not conducted. All procedures were reviewed and approved by the Chancellor’s Animal Research Committee (ARC) at the University of California, Los Angeles.

### Food restriction

One day before behavioral testing began, rats were singly-housed, the amount of chow given to each rat was reduced to 12 g/d, and rats were given ∼10 sucrose pellets (45-mg dustless precision sucrose pellets; Bio-Serv, Frenchtown NJ) in their home cage to acclimate them to the food rewards. Rats were maintained on 12g of food daily and were each fed within 30 min of completing the daily testing. Once rats progressed to the choice task they were given 8 g of chow per d, in addition to the food they consumed during testing. At the time of euthanasia, rats weighed an average of 356.5 g.

### Stereotaxic Surgery

General surgical procedures were the same as those recently published (Hart and Izquierdo, 2017). Rats were anesthetized with isoflurane (5% induction, 2% maintenance in 2L/min O_2_). Burr holes were drilled bilaterally on the skull for insertion of 26-gauge guide cannulae (PlasticsOne, Roanoke, VA), after which 33-gauge internal cannulae (PlasticsOne, Roanoke, VA) were inserted. Rats were infused with 0.5 μL of virus at a flow rate of 0.1 μL/minute and injectors were subsequently left in place for 5 additional minutes to allow for diffusion of solution. In the G_i_ experiment, the virus used was AAV8-CaMKIIα-hM4D(G_i_)-mCherry (Addgene viral prep # 50477-AAV8, Addgene, Cambridge, MA). In the G_q_ experiment, the virus used was AAV8-CaMKIIα-hM3D(G_q_)-mCherry (Addgene viral prep # 50476-AAV8, Addgene, Cambridge, MA). In the GFP (null virus) control, the virus used was AAV8-CaMKIIα-eGFP (Addgene, Cambridge, MA). In the imaging experiment, the virus used was AAV9-CaMKIIα-GCaMP6f (Addgene, Cambridge, MA). The coordinates used for the guide cannulae targeting ACC (Cg1) in the G_i_, G_q_, and GFP control experiments were: AP= +2.0 mm, ML= ±0.7 mm, DV= −1.9 mm from Bregma. Four of twelve rats in the G_i_ experiment received infusions in more anterior Cg1 to compare with other laboratory experiments on decision confidence (Stolyarova et al., 2019), at AP= +3.7 mm, ML= ±0.8 mm, DV= −1.6 mm from Bregma. Since no differences emerged from this differential targeting, we combined the Cg1 groups. In the imaging experiment, coordinates were: AP= +2.0 mm, ML= ±0.7 mm, DV= −1.4 mm (0.5 μL) from Bregma, and a second 0.5 μL bolus of virus was injected at DV= -0.9 mm. Injectors extended 1 mm beyond the tip of the cannula. Following the 5-minute diffusion time, the cannulae and injectors were removed, incisions were stapled closed, and the rats were placed on a heating pad and kept in recovery until ambulatory before being returned to the vivarium.

Three days following viral infusions in the subset of rats receiving AAV9-CaMKIIα-GCaMP6f (for calcium imaging), rats were implanted with 1.8 mm diameter 0.25 pitch GRIN lenses (Edmund optics part 64-519, Barrington, NJ). Following similar surgical procedures, 4 anchor screws were secured to the skull, after which a 2.0 mm craniotomy was drilled 0.2 mm lateral to the center of the viral infusion hole. The dura was cleared and approximately 0.5 mm of tissue was aspirated, after which the lens was placed 2.0 mm ventral from the surface of the skull and secured in place with cyanoacrylate glue and bone cement. The lens was protected with Kwik-Sil (World Precision Instruments, Sarasota, FL).

Post-operative care for all rats consisted of five daily injections of carprofen (5mg/kg, s.c.) and oral sulfamethoxazole/trimethoprim solution. Two-to-three weeks following lens implantation, a small aluminum baseplate was attached to the animal’s head and secured with bone cement. The exposed lens was cleaned with 100% ethanol, and the baseplate was secured in a position where cells in the field of view and vasculature were in focus. A 3D printed cover was secured to the baseplate with an anchor screw at all times when recording was not occurring. Rats were allowed a 5-d free feeding recovery period following viral infusion (G_i_, G_q_, and GFP experiments) or GRIN lens implantation (imaging experiment) after which they were food restricted and behavioral testing began.

### Apparatus

All behavioral testing was conducted in chambers outfitted with a house light, internal stimulus lights, a food-delivery magazine, and 2 retractable levers positioned to the left and right of the chamber wall, opposite the magazine. All hardware was controlled by a PC running Med-PC IV (Med-Associates, St. Albans, VT).

### Miniaturized microscope data collection

Microscopes were custom built according to plans available at Miniscope.org. Images were acquired with a CMOS imaging sensor (Labmaker, Berlin, Germany) attached to custom data acquisition (DAQ) electronics via a 1.5 mm coaxial cable. Data were transferred to a PC running custom written DAQ software over Super Speed USB. DAQ software was written in C++ and used Open Computer Vision (OpenCV) for image acquisition. 480×752 pixel images were acquired at 30 Hz and written to .avi files. DAQ software simultaneously recorded and time-stamped behavioral data and image data, allowing for offline alignment. All hardware design files and assembly instructions are available at Miniscope.org. Calcium signals were extracted using modified constrained non-negative matrix factorization scripts in Matlab (Mathworks 2016, Natick, MA) (Pnevmatikakis et al., 2016; Zhou et al., 2018).

### Lever press training

Rats were first given fixed-ratio (FR)-1 training where each lever press earned a single sucrose pellet (Bioserv, Frenchtown, NJ). They were kept on this schedule until they earned at least 30 pellets within 30 minutes. Following this, rats where shifted to a progressive ratio (PR) schedule where the required number of presses for each pellet increased according to the formula:

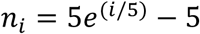

where *n*_*i*_ is equal to the number of presses required on the i^th^ ratio, rounded to the nearest whole number (Richardson and Roberts, 1996), after 5 successive schedule completions. No timeout was imposed. Rats were tested on the PR schedule until they earned at least 30 pellets on any given day (∼5 d) within 30 min. Upon meeting the PR performance criterion, a ceramic ramekin containing 18 g of lab chow was introduced (modified from (Randall et al., 2012)) during testing. Rats were free to choose between consuming freely-available but less preferred chow or lever pressing for preferred sucrose pellets. Rats (G_i_, G_q_, and GFP experiments) were given at least 5 choice testing sessions before clozapine-N-oxide (CNO) or vehicle (VEH) injections began (details below).

### Drug treatment during different types of test sessions

Rats in the G_i_, G_q_, and GFP control experiments were given either VEH (95% saline, 5% DMSO, 1 mL/kg) or CNO (3.0 mg/kg i.p. in 95% saline, 5% DMSO, 1 mL/kg) (Tocris, Bristol, UK) in counterbalanced order, 45 min prior to test sessions. Three types of test sessions were given: 1) a progressive ratio choice (PRC) session during which rats lever pressed on the PR schedule in the presence of the ceramic ramekin containing lab chow so that they were free to choose between lever pressing for sucrose versus free feeding on chow, 2) progressive ratio (PR) only sessions during which we omitted the ceramic ramekin (so that there was no freely-available lab chow) to assess whether manipulations decreased lever pressing in the absence of choice, and 3) a free choice consumption test where there was free access to pre-weighed amounts of sucrose pellets and lab chow (18 g) in empty cages (different from their home cages). Following this, any remaining food was collected and weighed to determine rats’ food preferences. All sessions were 30 min in duration. In a repeated-measures design, VEH or CNO was administered prior to a PRC testing session, a PR only testing session, and a free availability choice testing session, in that order, for each rat. The order of vehicle versus CNO administration was counterbalanced for baseline choice performance. Rats were given at least 48 hours between injections, and testing never occurred on consecutive days.

### Euthanasia

Following behavioral testing, rats were sacrificed by sodium pentobarbital overdose (Euthasol, 0.8 mL, 390 mg/mL pentobarbital, 50 mg/mL phenytoin; Virbac, Fort Worth, TX) and perfused transcardially with 0.9% saline followed by 10% buffered formalin acetate. Brains were post-fixed in 10% buffered formalin acetate for 24 hours followed by 30% sucrose cryoprotection for 5 days. 50 μm sections were cover slipped with DAPI mounting medium (Prolong gold, Invitrogen, Carlsbad, CA), and visualized using a BZ-X710 microscope (Keyence, Itasca, IL).

### DREADDs quantification

Fifty μm sections taken from each animal were visualized at seven AP coordinates relative to bregma: +4.2 mm, +3.7 mm, +3.2 mm, +2.7 mm, +2.2 mm, +1.7 mm, and +1.6 mm. No fluorescence was observed beyond these coordinates. mCherry fluorescence was drawn by a blind experimenter on a GNU Image Manipulation Program (GIMP) document containing a schematic of each of these seven sections, drawn to scale. Spread was quantified as total pixel count across all seven sections.

### Electrophysiological confirmation of DREADDs

Separate rats were prepared with ACC (Cg1) DREADDs using identical surgical procedures to the main experiments. Slice recordings did not begin until at least four weeks following surgery to allow sufficient hM receptor expression. Slice recording methods were similar to those previously published (Babiec et al., 2017). Six rats were deeply anesthetized with isoflurane and decapitated. The brain was rapidly removed and submerged in ice-cold, oxygenated (95% O_2_/5% CO_2_) artificial cerebrospinal fluid (ACSF) containing (in mM) as follows: 124 NaCl, 4 KCl, 25 NaHCO_3_, 1 NaH_2_PO_4_, 2 CaCl_2_, 1.2 MgSO_4_, and 10 glucose (Sigma-Aldrich). 400-μm-thick slices containing the ACC were then cut using a Campden 7000SMZ-2 vibratome. Slices from the site of viral infusion were used for validation. Expression of mCherry was confirmed after recordings were performed, and ACC slices with no transfection were used as control slices. Slices were maintained (at 30°C) in interface-type chambers that were continuously perfused (2–3 ml/min) with ACSF and allowed to recover for at least 2 hours before recordings. Following recovery, slices were perfused in a submerged-slice recording chamber (2–3 ml/min) with ACSF containing 100 μM picrotoxin to block GABA_A_ receptor-mediated inhibitory synaptic currents. A glass microelectrode filled with ACSF (resistance = 5–10 MΩ) was placed in layer 2/3 ACC to record field excitatory postsynaptic synaptic potentials and postsynaptic responses elicited by layer 1 stimulation delivered using a bipolar, nichrome-wire stimulating electrode placed near the medial wall in ACC. Inhibitory validation in ACC with identical coordinates, reagents, and virus was previously performed by our lab, with the methods and data appearing elsewhere (Stolyarova et al., 2019) and so these experiments were not needlessly repeated. Briefly, we first recorded for 2 minutes without synaptic stimulation to measure spontaneous levels of activity. Presynaptic fiber stimulation (0.2 msec duration pulses delivered at 0.33 Hz) was then delivered and the stimulation intensity was varied in 0.2 V increments to generate an input/output curve and identify the threshold for generation of postsynaptic responses. Stimulation strength was then set to the minimum level required to induce postsynaptic responses in ACC. Once stable responses (measured as the area of responses over a 4 second interval) were detected, baseline measures were taken for at least 10 minutes, followed by 20 minutes bath application of 10 μM CNO. In slices where CNO failed to elicit spontaneous activity, we generated a second input/output in the presence of CNO to test for CNO-induced changes in postsynaptic responses evoked by synaptic stimulation. Unless noted otherwise, all chemicals were obtained from Sigma-Aldrich.

### Behavioral analyses

Behavioral data were analyzed using GraphPad Prism v.7 (La Jolla, CA), SPSS v.25 (Armonk, NY), and MATLAB (MathWorks, Natick, Massachusetts; Version R2017a). An alpha level for significance was set to 0.05. A mixed ANOVA with between-subject factor of virus (G_i_, G_q_, eGFP null) and within-subject factor of effort condition and injection (PRC, PR; VEH, CNO) was conducted on lever pressing data as well as highest ratio achieved and number of pellets earned. Subsequently, paired samples t-tests (reported as means ± SEM) on data from the PRC and PR tests were used to test for effects of CNO vs. vehicle (VEH). Because the dependent measure was different in the free choice test (i.e. amount of food consumed, not lever presses), a separate mixed ANOVA with between-subject factor of virus (G_i_, G_q_, eGFP null) and within-subject factors of food type and injection (sucrose, chow; VEH, CNO) was conducted. Following these group comparisons which included virus as a factor, paired t-tests were used to test for effects of CNO on the total number of lever presses, highest ratio, and number of pellets earned in each of the groups. Two-way ANOVA was used to analyze the effects of CNO on temporal response patterns in each of the groups. Two-way ANOVA was used to test for effects of CNO on total consumption during free consumption testing in each of the groups. Mixed ANOVA was used to compare responding in the imaging and DREADDs animals, and repeated-measures ANOVA was used to compare responding in the different session types within the imaging group.

### Calcium image analyses

Image analyses were performed using custom written MATLAB (MathWorks, Natick, Massachusetts; Version R2017a) scripts. First, images were motion corrected using functions based on the Non-Rigid Motion Correction (“NoRMCorre”) package (Pnevmatikakis and Giovannucci, 2017), downsampled spatially by a factor of two (240×356 pixels) and temporally by a factor of 4 (to a frame rate of 7.5 fps). In order to remove background fluorescence, we performed a neural enhancement processing step as in the min1pipe processing framework (Lu et al., 2018). Briefly, from each frame we constructed a background fluorescence estimate by performing a neuron-sized morphological opening function, and subtracting this background frame from the original frame. This removes large fluorescence artifacts inherent in single photon microscopy, while preserving neural components. This motion corrected, downsampled, and enhanced video was then processed using Constrained Non-negative Matrix Factorization for Endoscopic data, “CNMF-E”, (Pnevmatikakis et al., 2016; Zhou et al., 2018). This extracted individual neural segments, denoised their fluorescent signals, demixed cross-talk from nearby neighbors, and deconvolved the calcium transients in order to estimate temporally constrained instances of calcium activity for each neuron (Friedrich et al., 2017). These estimated calcium event timings were used to compare calcium activity time-locked to specific behavioral instances. Neurons were matched across recording sessions using CellReg (Sheintuch et al., 2017) by matching cells based on their contours and centroid locations.

### Calcium response analysis

A cell’s probability of generating calcium transients proximal to a trigger event (such as a lever press, LP or magazine head entry, HE) was compared against a baseline probability surrounding each event. LP bouts were defined by the first lever press within a ratio and HE bouts defined as the first time point where the infrared beam was broken via nosepoke, after completion of that ratio. LP and HE bouts where the last lever press and first head entry bout were separated by more than 30 seconds were excluded from analysis. For every cell recorded within a session, we created a 6.25 s (±3.125) peri-event time histogram (PETH) divided into 47 equal time bins (bin size = 133 ms), with the 24^th^ (middle) bin centered upon the trigger events (which were either LP or HE bout starts). PETHs were constructed for LP and HE bouts from each sessions type, yielding 6 PETHs for each cell of the average transient rate surrounding each trigger event. Only half of the bout-centered PETH window (24 bins) was used as the ROI for statistical analysis.

To test whether a cell responded prior to LP events, we used a binomial test on each of the first 24 bins compared to the average transient rate in the 23 bins following LPs. The same procedure was applied to HE events, where a binomial test was applied to each of the last 24 bins compared to the average transient rate in the 23 bins preceding HEs. This pre-post event approach is consistent with what others have defined as task-evoked calcium responses (Jennings et al., 2015). These comparisons were used to identify cells modulated by each event while minimizing overlap between LP and HE responses (**Extended data 8-1**). A cell was classified as significantly responsive within the ROI if the transient rate within the ROI was greater than baseline, and one or more of three probability criteria were met: at least 1 out of 24 ROI bins beat p<.00125, at least 2 out of 24 ROI bins beat p<.0115, or at least 3 out of 24 ROI bins beat p<.0285. These p thresholds were chosen to yield an equal probability of each criterion being met, and p=.01 for meeting one or more of the criteria by chance. Hence, the probability of erroneously classifying a cell as responsive was 1%. Examples of LP and HE responsive cells are show in **Figure 7A** and **Figure 8A**, respectively.

To compare responses of each individual cell to LP and HE events during different types of experimental sessions, we analyzed a subpopulation of cells that met two criteria: 1) the cell responded significantly to the trigger event (LP or HE) during at least one session, and 2) the cell fired (but did not necessarily respond to the trigger event) during at least one session of all three imaging session types (PR, PRC, and CON). For each cell meeting these two criteria for LP events, three PETHs were derived, one for each session type: PETH_*PR*_, PETH_*PRC*_, PETH_*CON*_ (**Figure 7B**). For cells meeting these criteria for HE events, an additional PETH was computed for ramekin entries during PRC sessions (PETH_*R-PRC*_) to analyze similarity of responses between sucrose and chow consumption (**Figure 8B**). However, ramekin HEs within a session were few compared to sucrose HEs, and thus the PETH_*R-PRC*_ is relatively under sampled. Each mean PETH was constructed by computing an unweighted average of session PETHs over all events from the same session type during which the cell was active. The averaged PETH was then smoothed by convolving it with a 5-bin gaussian function of unit area. The smoothed PETH for each cell was normalized via division by a factor *B*, which was the value of the largest bin in any of the PETHs for that cell: *B* = max (PETH_*PR*_, PETH_*PRC*_, PETH_*CON*_). Finally, the frequency of responsive cells across session types for HE and LE events were compared against random number samples from a uniform distribution, analyzed using Chi-square tests.

### Satiety Control Condition

For a subset of calcium imaging sessions, we administered a satiety control condition, “CON.” For these sessions we capitalized on the fact that rats typically consume chow early in the test session, presented them with the lever and the chow initially, allowed them to consume chow, but then removed the chow once the rat began to lever press. This control condition allowed rats to reach a comparable motivational (more sated) state relative to the PR-only condition, and thus controlled for satiety differences between PR and PRC sessions. Lever pressing behavior in these sessions was similar to during PR-only sessions (**Extended data 7-1B**). The ramekin was removed after lever-pressing behavior commenced, so we could not construct ramekin response PETHs (PETH_*R-CON*_) since no ramekin HEs occurred after the 5^th^ lever press (defined by our baseline calcium transient rate calculation).

## RESULTS

### Chemogenetic manipulations

#### Histology

Reconstructions of viral spread confirmed that most placements were centered on the targeted region of Cg1 in ACC. Results of histological processing are shown in **Figure 1**. DREADD expression was driven by a CaMKIIα promoter, which is thought to selectively target projection neurons in cortex (Nathanson et al., 2009; Wang et al., 2013). Viral spread in the G_i_ and G_q_ groups was quantified by pixel count using GIMP software (see Methods). There was no significant difference between G_i_ and G_q_ groups (*t*_(20)_=2.03 *p*=0.056; G_i_ = 25258 ± 3121 total mean pixels; G_q_ = 17736 ± 2002 total mean pixels).

**Figure 1.**
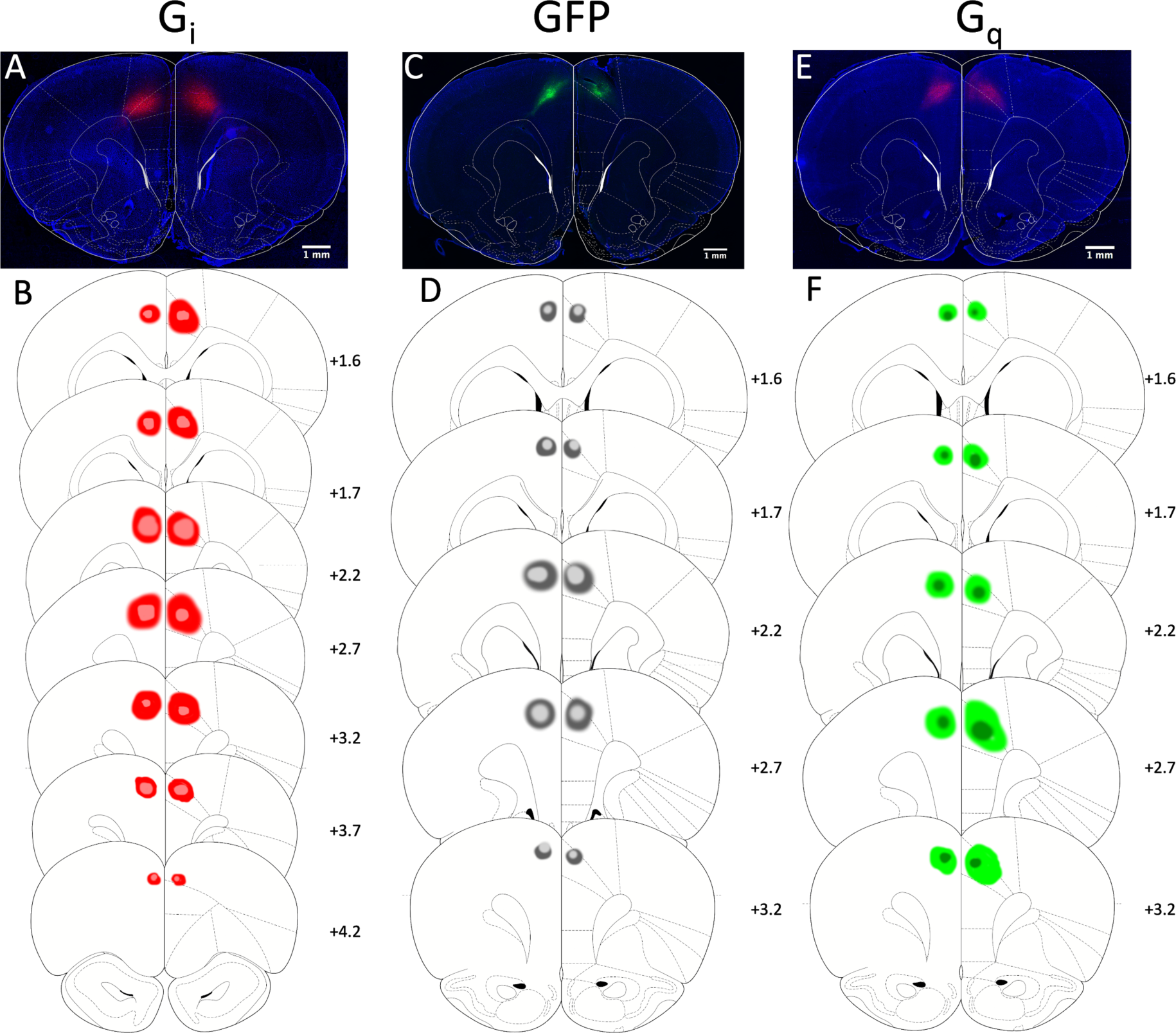
Expression of inhibitory and excitatory DREADDs and null virus eGFP in Anterior Cingulate Cortex. **(A)** Representative photomicrograph showing hM4Di-mCherry DREADDs under CaMKIIα in ACC, labeled G_i_. **(B)** Schematic reconstruction of maximum (red) and minimum (pink) viral spread for all rats. Numerals depict +Anterior-Posterior (AP) level relative to Bregma. Scale bars 1mm. **(C)** Representative photomicrograph showing eGFP (null virus) under CaMKIIα in ACC, labeled GFP. **(D)** Schematic reconstruction of maximum (dark grey) and minimum (light grey) viral spread for all rats. Numerals depict +Anterior-Posterior (AP) level relative to Bregma. Scale bars 1mm. **(E)** Representative photomicrograph showing hM3Dq-mCherry DREADDs expression in ACC, labeled G_q_. **(F)** Schematic reconstruction of maximum (light green) and minimum (dark green) viral spread for all rats. Numerals depict +Anterior-Posterior (AP) level relative to Bregma. Scale bars 1mm.

#### Ex vivo electrophysiological validation of DREADDs

We confirmed the efficacy of our DREADDs in slice recordings. A separate group of rats was prepared with G_q_ DREADDs in ACC (Cg1) using identical surgical procedures to the main experiments (**Figure 2A**). As described in Methods, a bipolar stimulating electrode was placed near the medial wall in layer I of ACC, and a glass microelectrode filled with ACSF (resistance = 5–10 MΩ) was placed in layer 2/3 of ACC to record field potentials and multiunit responses elicited by layer 1 stimulation. Using similar methods, we have previously reported that application of CNO strongly suppressed evoked field potentials in G_i_ transfected slices in ACC (Stolyarova et al., 2019). Here, we found that application of CNO induced spontaneous bursting in four out of six G_q_ transfected slices (spontaneous activity rate was increased from 0 to 0.06 Hz ± 0.01 Hz, inter-burst intervals were 18.3 ± 2.6 seconds (mean ± SEM, range: 12.4 – 31.6 seconds, n=4 slices from 3 rats) (**Figure 2B)**. In two other G_q_ transfected slices where no spontaneous bursting was observed, CNO application reduced threshold for stimulation induced postsynaptic responses (n=2 slices from 3 rats), with a sharp transition from no response to maximal response as a function of stimulation intensity (**Figure 2C)**. CNO had no effect on responses in non-transfected slices (n=3 slices from 3 rats) (**Figure 2D)**.

**Figure 2.**
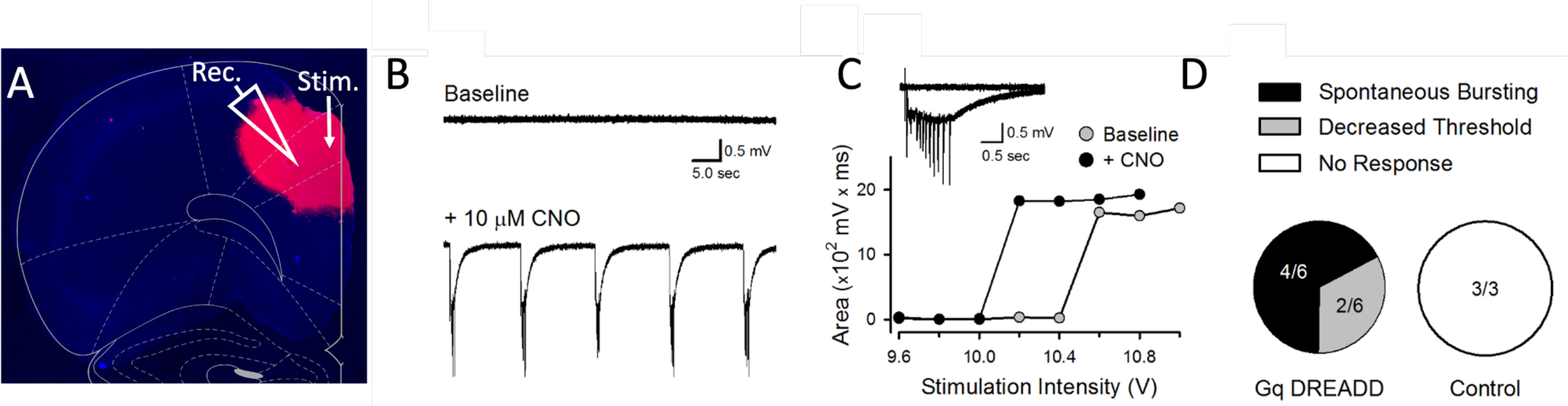
Excitatory hM3Dq DREADDs validation in Anterior Cingulate Cortex. Electrophysiological effect of bath application of CNO in slice. **(A)** Representative photomicrograph showing hM3Dq-mCherry DREADDs under CaMKIIα in a recording site of a 400 µm slice in ACC. **(B)** Spontaneous ACC activity in a G_q_ DREADD-expressing slice before (top) and after 20 min bath application of 10 mM CNO (bottom). CNO-induced spontaneous bursting was seen in 4 out of 6 G_q_ DREADD-expressing slices from 3 rats. **(C)** In the two G_q_ DREADD-expressing slices that failed to show spontaneous bursting there was a decrease in the threshold for stimulation induced postsynaptic responses. Plot shows results from one of these slices before (baseline) and after CNO application. **(D)** Summary of responses to CNO in G_q_ DREADD-expressing and control slices. CNO did not elicit either spontaneous bursting or decrease threshold for evoked responses in all 3 of the control (non-transfected) slices from 3 rats.

#### Group comparisons

Overall responding as measured by total lever presses was higher in PR than PRC sessions. A mixed ANOVA yielded a significant main effect of effort condition (F_(1,29)_=173.31 *p*<0.001). There was no main effect of virus type (G_i_, G_q_, eGFP null, F_(2,29)_=0.96 *p*=0.39). The interaction between effort condition (PRC, PR) and injection (VEH, CNO) approached the threshold for significance (F_(1,29)_=3.53 *p*=0.07). There was no significant group by effort condition by injection interaction (F_(2,29)_=0.74 *p*=0.48).

Identical analyses were performed for other measures of progressive ratio responding (highest ratio and number of pellets earned) with near identical patterns of results. As expected, animals achieved higher ratios in PR than PRC sessions, as revealed by a significant main effect of effort condition (F_(1,29)_=222.85 *p*<0.001). Highest ratio achieved was not different among the three groups; there was no main effect of virus type (G_i_, G_q_, eGFP null, F_(2,29)_=1.57 *p*=0.23). The interaction between effort condition (PRC, PR) and injection (VEH, CNO) on highest ratio approached the threshold for significance (F_(1,29)_=4.16 *p*=0.05). There was no significant group by effort condition by injection interaction (F_(2,29)_=0.63 *p*=0.54). Complementary to this, animals earned more pellets in PR than PRC sessions, as revealed by a main effect of effort condition (F_(1,29)_=238.33 *p*<0.001). The number of pellets earned was not different among the three groups; there was no main effect of virus (G_i_, G_q_, eGFP null, F_(2,29)_=0.92 *p*=0.41). There was a significant interaction between effort condition (PRC, PR) and injection (VEH, CNO) on number of pellets earned (F_(1,29)_=5.40 *p*=0.027). There was no significant group by effort condition by injection interaction (F_(2,29)_=0.55 *p*=0.58). Thus, virus groups did not differ by any measure of progressive ratio responding, though the effect of CNO, as tested by all three measures, did depend on which task animals were tested in – PR or PRC. To further clarify the effects of CNO, and test whether administration differentially affected PR or PRC testing, we conducted planned within-group comparisons, described in the next section.

Free choice consumption was not affected by CNO administration. There was no main effect of CNO (F_(1,29)_=0.21 *p*=0.65), nor was there a significant virus by injection (F_(2,29)_=0.43 *p*=0.65), virus by food type (F_(2,29)_=0.19 *p*=0.83), injection by food type (F_(1,29)_=0.33 *p*=0.57), or virus by injection by food type (F_(2,29)_=2.03 *p*=0.15) interaction. All animals consumed more sucrose than chow during free choice consumption, irrespective of virus or injection (F_(1,29)_=80.68 *p*<0.001). To more fully explore the effect of inhibition and excitation of ACC (i.e. the effects of CNO vs. VEH) on these measures, we conducted further analyses for each viral group, below.

#### Effects of DREADDs inhibition and excitation of ACC

We found that chemogenetic inhibition of ACC significantly decreased lever pressing during PRC but not PR sessions. Planned comparisons revealed CNO significantly reduced the total mean number of lever presses during 30 min PRC sessions (*t*_(10)_=3.047 *p*=0.01; VEH = 216.1 ± 52.85 presses; CNO = 173.2 ± 40.27 presses) **(Figure 3A)**. The total amount of chow consumed during 30 min PRC sessions was not affected by CNO treatment (*t*_(10)_=1.048 *p*=0.32; VEH = 7.79 ± 0.33 g; CNO = 7.50 ± 0.36 g) **(Figure 3B)**. CNO had no effect on PR responding in the absence of freely available chow (*t*_(10)_=0.26 *p*=0.80; VEH = 905.1 ± 102.00 presses; CNO = 893.9 ± 128.3 presses) **(Figure 3C)**.

**Figure 3.**
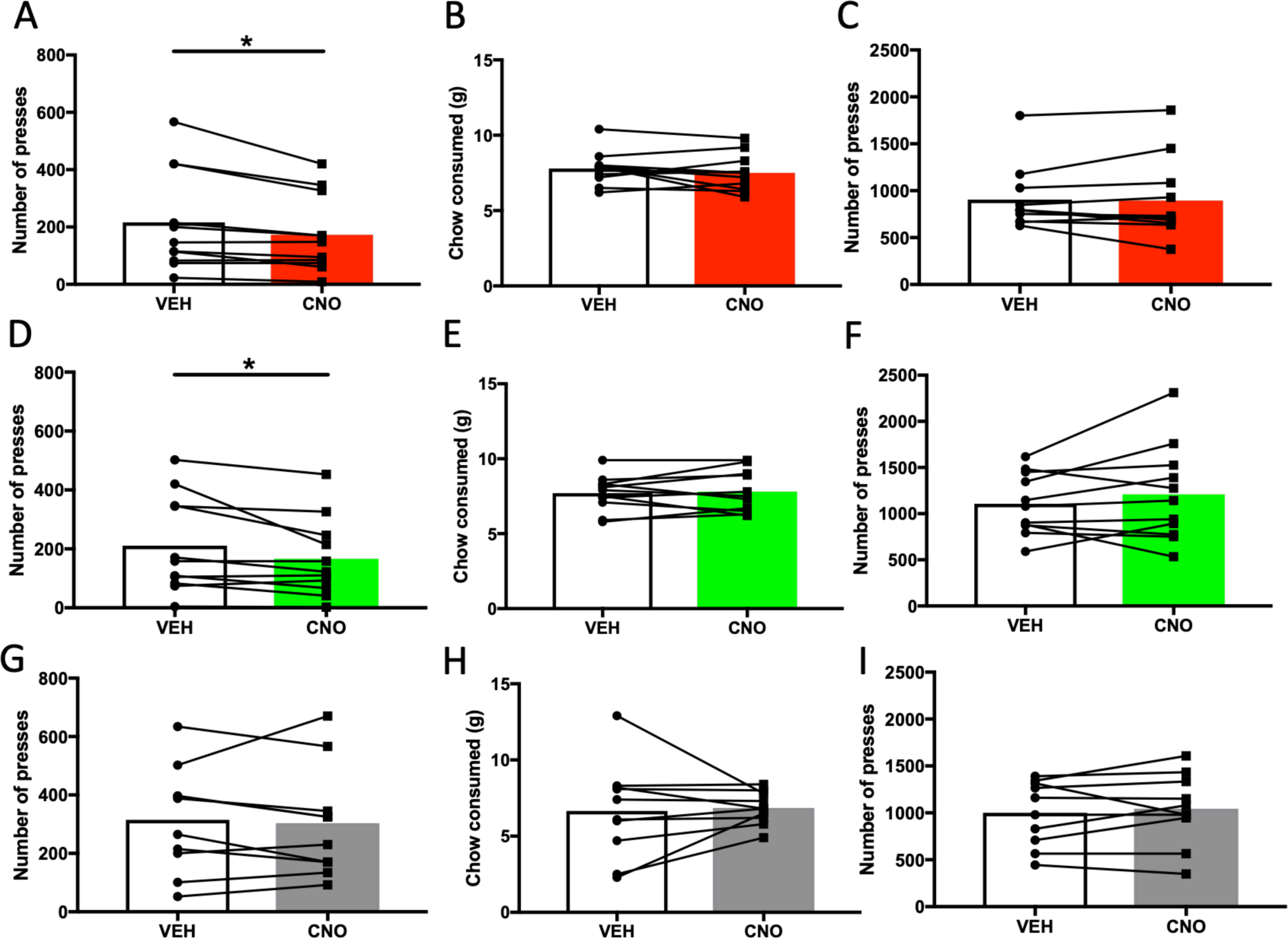
Effects on effortful choice behavior following DREADDs inhibition and excitation of Anterior Cingulate Cortex. **(A)** Mean lever presses during PRC sessions, when rats were presented with both the possibility of lever pressing under a PR schedule for sucrose pellets and freely-available chow. Shown is within-subject, counterbalanced choice behavior under vehicle (VEH) and CNO. CNO significantly reduced the number of lever presses in the G_i_ condition. **(B)** Total chow consumed during choice testing was not different following VEH vs. CNO in the G_i_ condition. **(C)** Mean lever presses during PR sessions when there was no chow available as an alternative option. Lever pressing was not different following VEH vs. CNO in the G_i_ condition. **(D)** CNO significantly reduced the number of lever presses for sucrose pellets in the G_q_ condition, when chow was also available. **(E)** Total chow consumed during choice testing was not different following VEH vs. CNO in the G_q_ condition. **(F)** Lever pressing was not different following VEH vs. CNO in the G_q_ condition, when chow was not available. **(G)** No effect on effortful choice behavior following CNO administration in GFP (null virus) controls when chow was also available. **(H)** Total chow consumed during choice testing was not different following VEH vs CNO in the GFP condition. **(I)** Lever pressing was not different following VEH vs. CNO in the GFP condition, when chow was not available. * *p*<0.05.

Using identical procedures to the inhibition experiment, a separate group of rats was assessed on effortful choice following excitatory G_q_ DREADDs transfection in ACC. Similar to results for the G_i_ experiment, we found that chemogenetic excitation of ACC significantly decreased lever pressing during PRC but not PR sessions. Planned comparisons revealed CNO significantly reduced the total mean number of lever presses during 30 min PRC sessions (*t*_(10)_=2.31 *p*=0.04; VEH = 210.7 ± 49.44 presses; CNO = 166.6 ± 40.4 presses) **(Figure 3D)**. The total amount of chow consumed during 30 min PRC sessions was not affected by CNO treatment (*t*_(10)_=0.39 *p*=0.71; VEH = 7.71 ± 0.36 g; CNO = 7.81 ± 0.42 g) **(Figure 3E)**. CNO had no effect on PR responding in the absence of freely available chow (*t*_(10)_=1.15 *p*=0.28; VEH = 1107.00 ± 99.66 presses; CNO = 1209.00 ± 156.3 presses) **(Figure 3F)**.

To test whether results for G_i_ and G_q_ DREADDs could be explained by virus exposure alone, a control virus carrying only GFP (but no DREADD receptors) was infused into ACC in a separate group of rats. A paired t-test revealed no effect of CNO on the total mean number of lever presses during 30 min PRC sessions (*t*_(9)_=0.48 *p*=0.64; VEH = 314.9 ± 57.34 presses; CNO = 302.7 ± 59.66 presses) **(Figure 3G)**. The total amount of chow consumed during 30 min PRC sessions was not affected by CNO treatment (*t*_(9)_=0.26 *p*=0.80; VEH = 6.65 ± 0.99 g; CNO = 6.85 ± 0.34 g) **(Figure 3H)**. CNO also had no effect on PR responding in the absence of freely available chow (*t*_(10)_=0.73 *p*=0.48; VEH = 1001.00 ± 109.4 presses; CNO = 1043.00 ± 120.00 presses) **(Figure 3I)**.

The same pattern was observed for conventional measures of PR responding, highest ratio and number of pellets earned, during PRC sessions (**Figure 3-1**). In the G_i_ group, CNO reduced the highest ratio achieved (*t*_(10)_=3.48 *p*=0.0059; VEH = 14.91 ± 2.63; CNO = 12.36 ± 2.04) and number of pellets earned (*t*_(10)_=4.209 *p*=0.0018; VEH = 30.18 ± 3.3 pellets; CNO = 27.45 ± 3.2 pellets). In the G_q_ group, CNO reduced the highest ratio achieved (*t*_(10)_=2.43 *p*=0.0355; VEH = 14.36 ± 2.68; CNO = 11.82 ± 2.05) and number of pellets earned (*t*_(10)_=2.70 *p*=0.022; VEH = 29.36 ± 3.71 pellets; CNO = 26.55 ± 3.50 pellets). CNO had no effect on the highest ratio achieved (*t*_(10)_=0.33 *p*=0.75; VEH = 19.9 ± 2.80; CNO = 20.5 ± 3.06) or the number of pellets earned (*t*_(10)_=0.29 *p*=0.78; VEH = 35.2 ± 3.69 pellets; CNO = 34.6 ± 2.54 pellets) in the GFP control group.

As with lever pressing, these measures were not affected during PR sessions. In the G_i_ group, CNO had no effect on highest ratio achieved (*t*_(10)_=0.71 *p*=0.49; VEH = 43.00 ± 3.89; CNO = 44.36 ± 4.47) or number of pellets earned (*t*_(10)_=0.07 *p*=0.95; VEH = 54.55 ± 1.88 pellets; CNO = 54.45 ± 2.43 pellets). In the G_q_ group, CNO had no effect on highest ratio achieved (*t*_(10)_=1.04 *p*=0.32; VEH = 54.91 ± 4.47; CNO = 58.82 ± 5.71) or number of pellets earned (*t*_(10)_=0.87 *p*=0.41; VEH = 59.09 ± 1.77 pellets; CNO = 60.45 ± 2.46 pellets). CNO also had no effect on the highest ratio achieved (*t*_(10)_=0.72 *p*=0.49; VEH = 49.7 ± 4.56; CNO = 51.5 ± 5.12) or the number of pellets earned (*t*_(10)_=0.60 *p*=0.56; VEH = 57.2 ± 2.38 pellets; CNO = 57.8 ± 2.59 pellets) in the GFP control group.

#### Free choice consumption tests

In the G_i_ group, a two-way ANOVA (food type-sucrose, chow; injection-VEH, CNO) on the amount of sucrose and chow consumed during free choice consumption testing revealed a significant main effect of food type (F_(1,10)_=12.02 *p*=0.006; sucrose = 8.59 ± 0.83 g; chow = 4.19 ± 0.34 g), but no significant food type by injection interaction (F_(1,10)_=1.415 *p*=0.26), or main effect of injection (F_(1,10)_=0.01 *p*=0.91) was found. Hence, food preference was intact and not affected by CNO injection. In the G_q_ group, a two-way ANOVA (food type-sucrose, chow; injection-VEH, CNO) on the amount of sucrose and chow consumed during free choice consumption testing revealed a significant main effect of food type (F_(1,10)_=47.63 *p*<0.001; sucrose = 9.68 ± 0.47 g; chow = 5.10 ± 0.37 g), but no significant food type by injection interaction (F_(1,10)_=2.27 *p*=0.16) or main effect of injection (F_(1,10)_=0.06 *p*=0.81), indicating food preference was intact and unaffected by CNO. Similarly, in the GFP group, a two-way ANOVA (food type-sucrose, chow; injection-VEH, CNO) on the amount of sucrose and chow consumed during free choice consumption testing revealed a significant main effect of food type (F_(1,9)_=80.63 *p*<0.001; sucrose = 8.37 ± 0.60 g; chow = 3.19 ± 0.35 g), indicating food preference was intact. No significant food type by injection interaction (F_(1,9)_=1.27 *p*=0.29), or effect of injection (F_(1,9)_=0.97 *p*=0.35) was found **(Figure 4)**.

**Figure 4.**
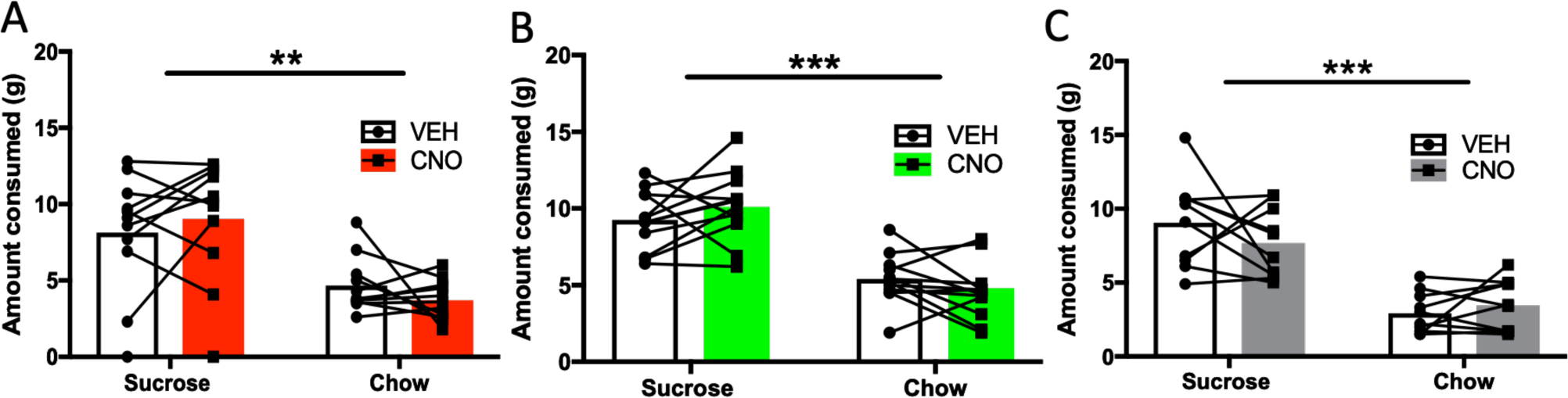
Free choice consumption in all three treatment groups following CNO administration. Mean consumption of sucrose and chow when rats were presented with both as freely-available options. Shown is within-subject, counterbalanced choice behavior under vehicle (VEH) and CNO, indicating that food preference was intact: rats preferred the sucrose over chow. **(A)** CNO had no effect on either sucrose or chow consumed in the G_i_ condition. **(B)** CNO had no effect on either sucrose or chow consumed in the G_q_ condition. **(C)** CNO had no effect on either sucrose or chow consumed in the null virus (GFP) condition. **p<0.01, ***p<0.001.

#### Timecourse of lever pressing in PR and PRC

We found that the presence of an alternative option (chow) reduced lever pressing in the PRC condition, and that both G_i_ or G_q_ DREADDs decreased lever pressing in this PRC condition, but not in the PR condition where lever pressing was more robust. Of course, reduced total lever pressing over the course of thirty minutes does not necessarily indicate that lever pressing was unaffected during PRC testing. It is possible that G_i_ and G_q_ transfected animals may have shown a different temporal pattern of lever pressing behavior, despite showing a similar number of total presses. Likewise, it is possible that CNO *did* affect responding during PR testing, by altering the time course of presses rather than total number of presses. We therefore assessed the timecourse of lever pressing in 5 min time bins, in PR and PRC session types, in G_i_ and G_q_ DREADDs, following CNO and vehicle (VEH) injections (**Figure 5**).

**Figure 5.**
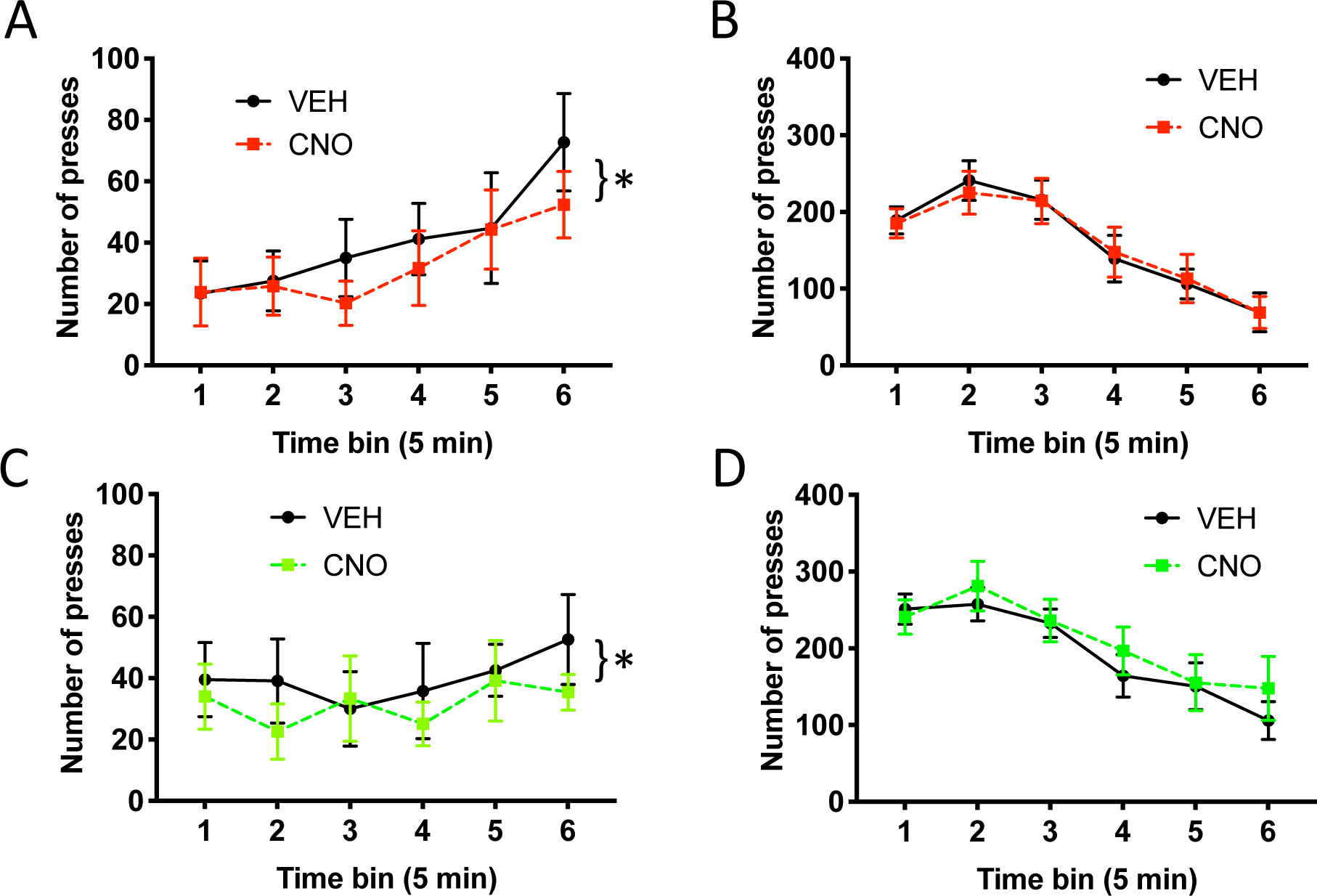
Time course of lever pressing in different session types. Mean lever pressing across 5 min time bins. **(A)** Lever pressing during PRC sessions, when rats were presented with both sucrose pellet and chow options in the G_i_ condition, in rats receiving CNO compared to within-subject vehicle (VEH). **(B)** Lever pressing during PR sessions, when rats were presented with a single option in the G_i_ condition, in rats receiving CNO compared to VEH. **(C)** Lever pressing in PRC sessions, when rats were presented with both options in the G_q_ condition, in rats receiving CNO compared to VEH. **(D)** Lever pressing during PR sessions, when rats were presented with a single option in the G_q_ condition, in rats receiving CNO compared to VEH. Bars represent S.E.M. *p<0.05.

As above, we first conducted a mixed ANOVA to test for group differences in temporal pressing patterns between the G_i_ and G_q_ DREADDs groups. Two separate mixed ANOVAs with virus as a between-subject factor, and injection and time bin as within-subject factors were conducted on PRC and PR data. We were unable to include the GFP group here because these data were lost due to hardware failure. Virus groups did not differ in the time course of lever pressing during PRC testing. A mixed ANOVA did not yield a significant main effect of virus on PRC pressing (F_(1,20)_=0.01 *p*=0.93). Responding increased across PRC sessions as revealed by a main effect of time bin (F_(5,100)_=3.17 *p*=0.01), and CNO generally suppressed lever pressing as revealed by a main effect of CNO (F_(1,20)_=14.30 *p*=0.001). There was no significant interaction between virus x injection (F_(1,20)_=0.02 *p*=0.88), virus x bin (F_(5,100)_=0.98 *p*=0.43), or virus x injection x bin (F_(5,100)_=0.72 *p*=0.61). Virus groups also did not differ in the time course of lever pressing during PR testing. A mixed ANOVA did not yield a significant main effect of virus on PR pressing (F_(1,20)_=2.11 *p*=0.16). Responding changed over the course of PR sessions as revealed by a main effect of time bin (F_(5,100)_=32.729 *p*<0.0001). In contrast to PRC testing, CNO had no effect on the pattern of PR lever pressing: there was no significant main effect of CNO (F_(1,20)_=0.84 *p*=0.37). There was no significant interaction between virus x injection (F_(1,20)_=1.08 *p*=0.31), virus x bin (F_(5,100)_=0.51 *p*=0.64), or virus x injection x bin (F_(5,100)_=0.42 *p*=0.83) during PR testing.

We separately tested the G_i_ and G_q_ DREADDs groups for effects of CNO. For the G_i_ DREADDs PRC condition, a two-way ANOVA revealed a significant effect of time bin (F_(5,50)_=3.423 *p*<0.01), a significant effect of injection type (F_(1,10)_=9.44 *p*<0.02; CNO = 33.06 ± 13.50 presses; VEH = 40.77 ± 16.65 presses, mean presses per time bin), but no significant time bin x injection type interaction (F_(5,50)_=0.88 *p*=0.50). For the G_i_ DREADDs PR condition, a two-way ANOVA resulted in a significant effect of time bin (F_(5,50)_=21.27 *p*<0.001), but no significant effect of injection type (F_(1,10)_=0.02 *p*=0.89) or time bin x injection type interaction (F_(5,50)_=0.15 *p*=0.98).

For the G_q_ DREADDs PRC condition, a two-way ANOVA revealed no significant effect of time bin (F_(5,50)_=0.577 *p*=0.71), a significant effect of injection type (F_(1,10)_=5.93 *p*=0.04; CNO = 31.62 ± 12.91 presses; VEH = 39.94 ± 16.31 presses, mean presses per time bin), but no significant interaction of time bin by injection type (F_(5,50)_=0.766 *p*=0.58). Finally, for the G_q_ DREADDs PR condition, a two-way ANOVA resulted in a significant effect of time bin (F_(5,50)_=13.03 *p*<0.001), but no significant effect of injection type (F_(1,10)_=1.20 *p*=0.30), or time bin by injection type interaction (F_(5,50)_=0.725 *p*=0.61). Thus, the reduction in total lever presses observed during PRC exhibited a similar pattern in G_i_ and G_q_ groups, and the pattern of responding was completely unaffected during PR testing. CNO only reduced responding during PRC testing, and it did so similarly in both DREADDs groups. We concluded that ACC interference disrupts high-effort lever pressing behavior when a choice is involved, but not when it is the only food option available.

### In-vivo calcium imaging

To further investigate why DREADD manipulations in ACC exerted their effects only in the choice condition, but not when PR lever pressing was the only food option, we performed in-vivo calcium imaging experiments to track responses of individual ACC cells and compare their activity during PR versus PRC sessions. Imaging rats were also given “satiety control” (CON) sessions, where rats were pre-fed with chow in the operant chamber, and then were allowed to lever press on a PR schedule in the absence of chow. This was to control for the possibility that neural encoding could be influenced by satiety from chow consumption, regardless of whether chow was freely available during lever pressing. Behavior during imaging sessions was similar to behavior in vehicle groups from the chemogenetics experiments (**Extended data 7-1A**), with the highest levels of responding occurring during the early portion of PR sessions (tapering off later in PR sessions), and steady low rates of responding throughout PRC sessions. A mixed ANOVA with experiment group (GCaMP, DREADDs) as a between-subject factor and session type (PR, PRC) and time bin (5 min) as within-subject factors revealed a significant main effect of time bin (F_(5,125)_=5.27 *p*=0.0002). Not surprisingly, overall response rates were lower in the imaging group than in the vehicle groups, as revealed by a significant main effect of experiment group (F_(1,113)_=21.04 *p*<0.0001) (**Extended data 7-1A**). This is due to rats wearing the miniscope and being tethered by the coaxial cable, which inhibited pressing behavior. But the effect of time on lever pressing behavior across PR and PRC sessions did not depend on experiment group-there was no significant interaction between time bin x experiment group (F_(5,113)_=1.10 *p*=0.37). A one way repeated-measures ANOVA was used to test for differences in the total number of lever presses, averaged across all imaging sessions, in the four rats included in our calcium analyses, during PR, PRC, and CON sessions. As expected, session type did affect the total number of presses as revealed by a significant main effect of session type (F_(2,6)_=9.13 *p*=0.02). Post-hoc comparisons using Tukey’s correction for familywise error revealed that the total number of presses was lower during PRC than PR sessions (*p*=0.04) as well as CON sessions (*p*=0.017). There was no significant difference between PR and CON sessions (*p*=0.79). Imaging rats showed reduced pressing during PRC sessions, much like vehicle rats (**Extended data 7-1B**).

#### Calcium imaging in rat ACC

A group of 4 rats received infusions of AAV9-CaMKIIα-GCaMP6f and was subsequently implanted with 1.8 mm diameter 0.25 pitch GRIN lenses in Cg1 (**Figure 6**). Rats were then trained on lever pressing and tested during PR and PRC sessions (identical to those described above for DREADD experiments), as well as CON sessions described above. We recorded a total of 1151 neurons from ACC from the 4 animals (136 from rat #1, 567 from rat #2, 254 from rat #3, 194 from rat #4). Each neuron’s calcium events were extracted by deconvolving its denoised calcium trace (sampling rate = 7.5 Hz; see Methods). To analyze neural responses to LP and HE events, perievent time histograms (PETHs) were generated by combining data from all sessions of a given type (PR, PRC, or CON) during which the cell exhibited at least one calcium event. Calcium event probabilities were computed in 133.33 ms time bins within ±3 s of the trigger event (LP or HE). To analyze how neural activity varied with session type (PR, PRC, CON), we identified a subset of 227 neurons (20% of all recorded cells) that were: 1) active during at least one session of each type (PR, PRC, and CON), and 2) significantly responsive to either LP or HE events (or both, see below for responsiveness criteria) in the session-averaged PETH for at least one of the three session types (**Figure 6H**). Cells (n=924) that did not meet these two criteria were excluded from further analyses.

**Figure 6.**
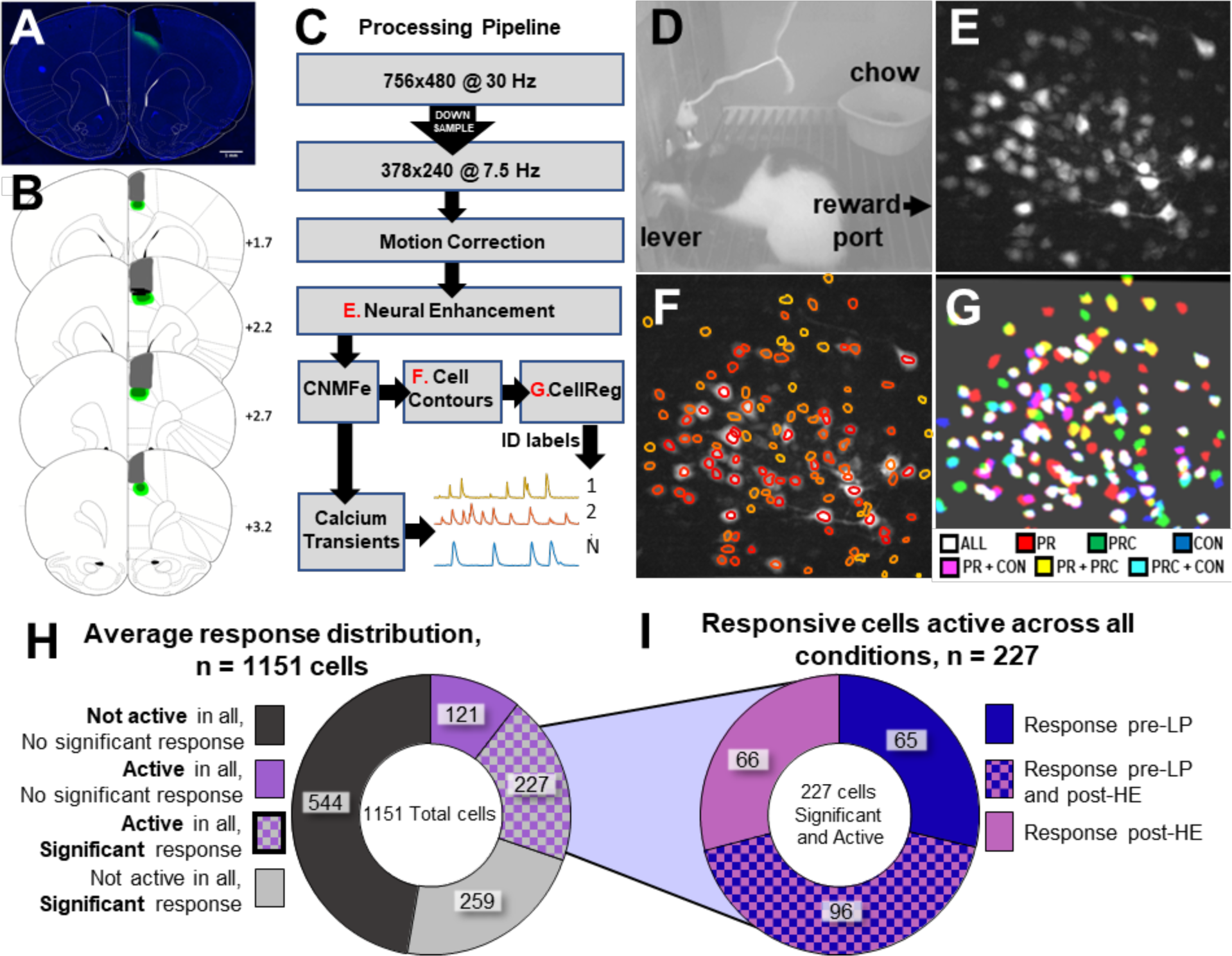
Calcium imaging during lever pressing. **(A)** Representative photomicrograph showing GCaMP6f expression and aspiration site for lens placement for ACC imaging. **(B)** Schematic reconstruction of maximum (light green), minimum (dark green) viral spread, and maximum aspiration damage (gray). Black bars represent ACC recording sites. Numerals depict +Anterior-Posterior (AP) level relative to Bregma. Scale bar 1mm. **(C)** Flow diagram for calcium imaging analysis pipeline. **(D)** Example of PRC behavior session with the miniscope on the rat’s head, and both sucrose and chow options available; reward port is on the right just out of view, chow is located in ramekin. **(E)** Max projection image from the session in D after motion correction and neural enhancement. **(F)** Same as in E, now with extracted cell contours overlaid. **(G)** Contours matched across three different recording sessions, each taken 6 days apart. **(H)** Proportions of cells that were active or responsive to stimuli. Cells that were active in at least one session of each type (PR, PRC, and CON) and had a significant response to either lever-press (LP) or head-entry (HE) events were included in further analysis. **(I)** Proportions of cells that were responsive to LP and HE events.

#### Responses preceding LP bouts

We analyzed neural responses occurring prior to the onset of each bout of lever pressing, under the assumption that this is a likely time window during which the decision to exert effort (that is, press the lever) is made. The onset of an LP bout was defined as the first LP that occurred after a magazine head entry and was followed by a HE within 30 seconds after completion of the lever-press ratio trial. LP response PETHs were triggered only by these LP onset events, not by all LP events (**Figure 7A**). A neuron was classified as LP-responsive if it showed a significantly higher probability of generating calcium events compared to the baseline rate within that session (see Methods). About 60% (96/161) of LP-responsive neurons were also responsive to magazine head entries (**Figure 6I**), and these were included in the analyses of LP-responsive cells presented below. These cells (significantly responsive to LPs and HEs) were typically due to significant responding on separate sessions.

**Figure 7.**
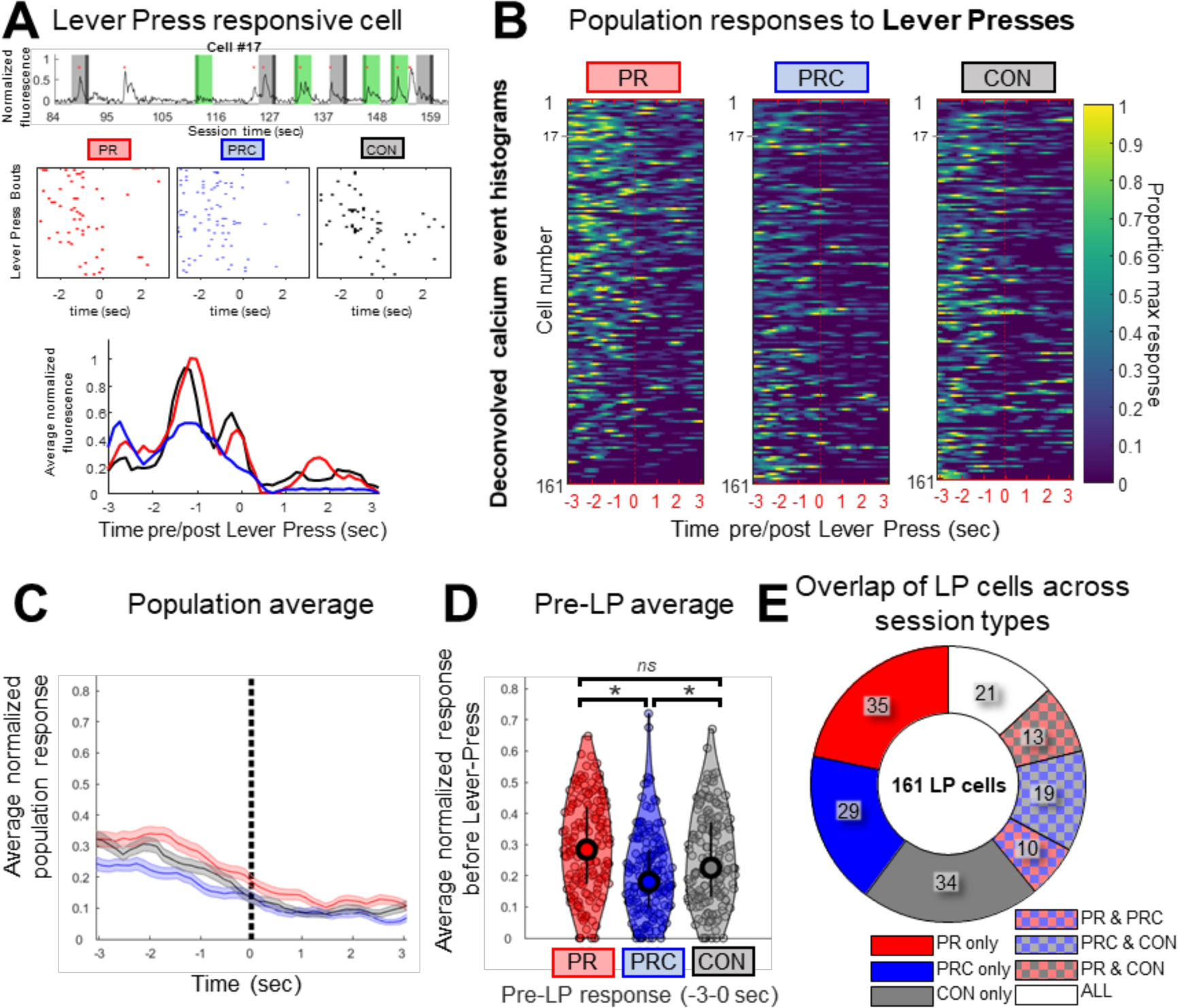
Neural responses in ACC prior to lever press bouts. **(A)** (Top) Example of raw fluorescence from an LP-resonsive cell during a PR session. Light grey bands indicate 3 second window before lever press bout begins, dark grey is the first lever press. Light green bands indicate 3 second window after the first head entry following the lever press bout; dark green is the first head entry timestamp. Red dots indicate times at which deconvolved calcium transients occurred. (Middle) Rastergrams of calcium events show that this cell often fired within ∼3 s prior to lever presses during PR, PRC, and CON sessions. (Bottom) Smoothed PETHs for this cell during each session type. **(B)** Heat maps show PETHs for all 161 cells that responded significantly prior to LP events. Color scale in each row is normalized to the max PETH bin value observed for the cell in that row across all three session types. **(C)** Mean of PETHs for each session type in B. **(D)** Mean normalized area under PETH curve for LP-responsive cells during the 3 s prior to LP events for PR, PRC, or CON sessions. **(E)** Proportions of LP-responsive cells that were significantly responsive during all 7 possible combinations of session types. Bars and shaded regions represent ±SEM, *p<0.05, *ns* = nonsignificant after accounting for multiple comparisons.

For every LP-responsive cell, three LP-triggered PETHs were generated (one for each session type: PR, PRC, CON; **Figure 7B**). Each PETH plotted the mean calcium event rate per time bin, averaged over all sessions of a given type during which the cell was active. To normalize the response of each cell across session types, the bins of all three PETHs for the cell were divided by the maximum value observed in any bin from all three PETHs (**Figure 7A, bottom**). A population averaged LP response curve was computed by taking the mean of the normalized PETHs for each session type (**Figure 7C**). To statistically compare the pre-LP responses of neurons in the population, we computed the mean normalized response 3 seconds before lever press bouts for LP responsive cells (**Figure 7D**). By this measure, it was found that pre-LP responses differed significantly in magnitude by session type (Friedman’s non-parametric ANOVA: *p* = 2.90e-5). Post-hoc comparisons revealed that the pre-LP response was significantly larger during PR and CON sessions compared to PRC sessions (Sign rank test: PR > PRC, *p* = 1.35e-05; PRC > CON, *p* = 0.0017), whereas PR and CON pre-LP responses were not significantly different after correcting for multiple comparisons (Sign rank test: PR = CON, *p* = .033). Hence, at the population level, ACC neurons were significantly less responsive before the onset of lever press bouts when the rat was offered the choice of free chow from the ramekin as an alternative to lever pressing during PRC sessions, and this effect could not be accounted for by satiety. We next evaluated how significantly responding cells were distributed across the different session types, and found that 60% (100/161 cells) were only responsive to LPs during one session type, which is not significantly different from what would be expected from a random allocation (expected: 102/161 cells; *Χ* ^2^ = 0.428, *p* = 0.52, **Figure 7E**). However, we found that 21 cells were significantly response to LPs during all three session types, greater than would be expected from chance (expected: 8/161 cells; *Χ* ^2^ = 22.23, *p* < 0.0001).

#### Responses following HE events

We analyzed neural responses occurring immediately after magazine HE events, under the assumption that this is a likely time window during which signals encoding the value of the sucrose reward relative to the free alterative (chow) might be generated; as the reward is experienced (**Figure 8A**). Only HE that occurred after completion of a lever pressing bout were included in the analysis, so HE events that occurred during the chow consumption period prior to lever pressing in PRC and CON sessions were not included. A neuron was classified as HE-responsive if it exhibited a significantly higher probability of generating calcium events during the 3 seconds after HE compared to the baseline rate within that session (see Methods). About 60% of HE-responsive neurons were also responsive to LP bout onset (**Figure 6I**), and these were included in the analyses of HE-responsive cells presented below.

**Figure 8.**
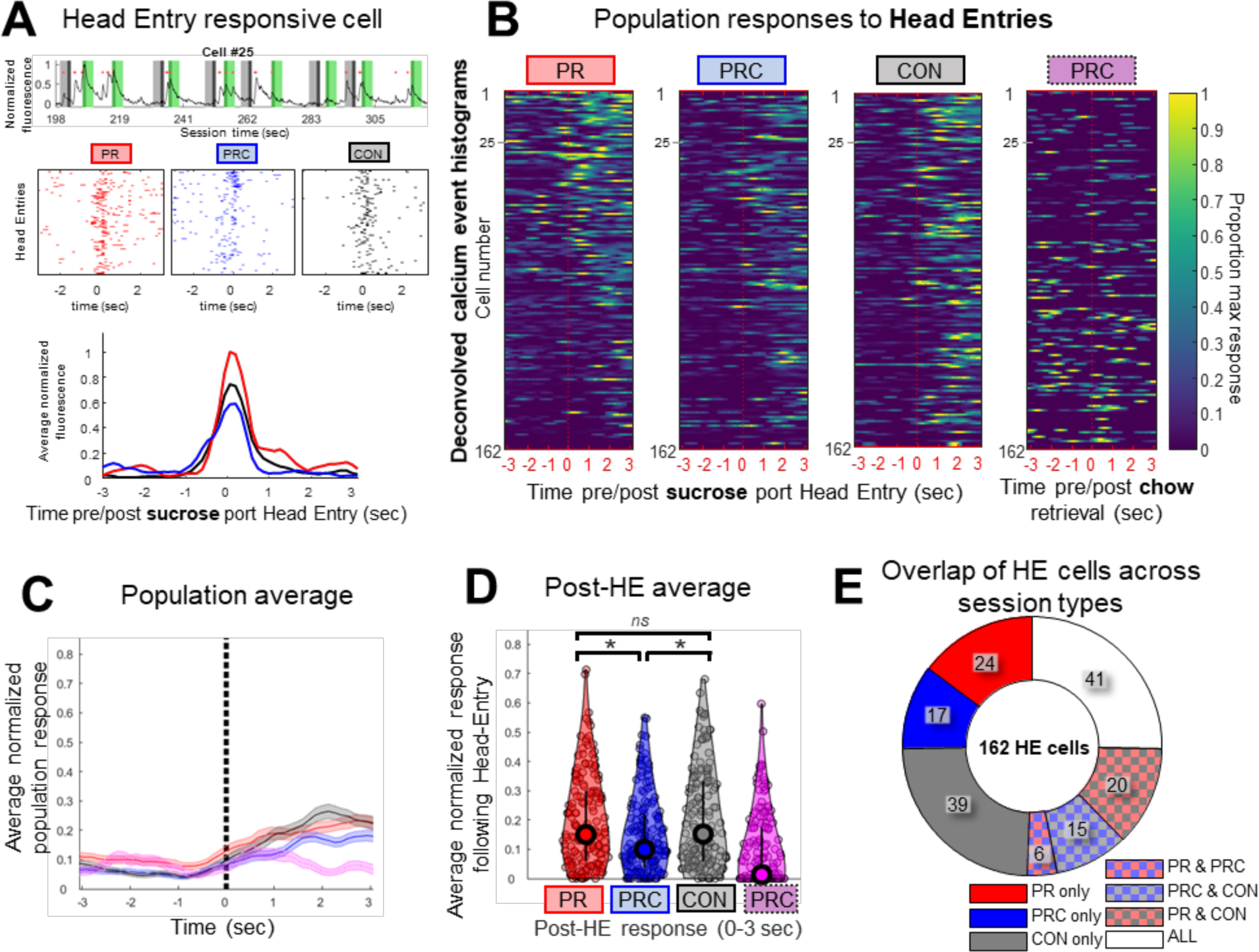
Neural responses in ACC following magazine head entry for sucrose reward. **(A)** (Top) Example of raw fluorescence from an HE-responsive cell during a PR session, plotted as in Figure 7A. (Middle) Rastergram of calcium event times show that this cell often fired just after HE events in PR, PRC, and CON sessions. (Bottom) Smoothed PETHs for this cell during each session type. **(B)** Heat maps show PETHs for all 162 cells that responded significantly to HE events. For comparison, PETHS triggered by ramekin entries during PRC sessions are also shown (right). Color scale in each row is normalized to the max PETH bin value observed for the cell in that row across PR, PRC, and CON sessions. **(C)** Mean of PETHs for each session type in B. **(D)** Mean normalized area under PETH curve for HE-responsive cells during the 3 s following HE events for PR, PRC, or CON sessions. **(E)** Proportions of HE-responsive cells that were significantly responsive during all 7 possible combinations of session types. Bars and shaded regions represent ±SEM, *p<0.05, *ns* = nonsignificant after accounting for multiple comparisons.

Three HE-triggered PETHs were generated for each cell (one for each session type: PR, PRC, CON), as well as an additional PETH triggered by head entries into the chow ramekin during PRC sessions (**Figure 8B**). Head-entries to the chow ramekin during the CON session were not analyzed, since these ramekin entries occurred prior to lever pressing, during the time period that was excluded from analysis of HE reponses. Population averaged responses to HE events were computed for each of the three session types (**Figure 8C**), using the same methods described above for LP-triggered PETHs. It was found that post-HE responses differed significantly by session type (Friedman’s non-parametric ANOVA: *p* = 0.046; **Figure 8D**). Post-hoc comparisons revealed that population responses during PR and CON sessions were not significantly different from one another (Sign rank test: PR = CON, *p* = 0.32), whereas PRC sessions showed a significantly lower post-HE response than either of the other two session types (Sign rank test: PR > PRC, *p* = 0.0021 ; CON > PRC, *p* = 1.28e-04). This pattern of results suggests that population averaged responses of ACC neurons to rewarding outcomes were smaller in the presence of available alternative outcomes than when no alternative outcome was available, and this effect could not be explained by satiety. Additionally, we found that the distribution of significantly responding HE cells was different than for LP cells (**Figure 8E**). More cells responded significantly to HEs in only a single session than would be expected from chance (80/162 cells, expected: 102/162; *Χ* ^2^ =12.812, *p* = 0.0003), half of which (49%) were cells only responding during the CON sessions. Of the remaining population responding significantly in more than one session, 41 HE cells were responsive during all three session types, also greater than expected from chance (41/162 cells, expected: 9/162; *Χ* ^2^ =120.47, *p* < 0.00001).

## DISCUSSION

We report that either chemogenetic silencing *or* stimulation of ACC excitatory neurons resulted in decreased effort for a qualitatively preferred option, and that this effect was only observed when a concurrently available, lower effort alternative was available, not when lever pressing was the only response option. Chemogenetic manipulations had no effect on the ability to lever press for sucrose or on food preference. CNO administration also had no effect in rats lacking active DREADD (hM4D-G_i_ or hM3D-G_q_) receptors. Slice electrophysiology confirmed robust inhibition (Stolyarova et al., 2019) and excitation in hM4D-G_i_ and hM3D-G_q_ transfected slices, respectively. Finally, using single-photon imaging, we found that ACC neurons showed differential task-evoked activity during lever pressing and reward-retrieval behavior that depended on the availability of another food option. In the same way that interference only affected choice lever-pressing, we found that tracked ACC neurons exhibited different response profiles during PR and PRC sessions. Taken together, these findings support a role for ACC in the evaluation of effortful behavior, consistent with recent evidence from single-unit recordings (Porter et al., 2019) and human fMRI (Arulpragasam et al., 2018).

### ACC chemogenetic silencing

The earliest studies probing rat ACC in effort-based choice made use of T-maze tasks where rats selected between the same food option but of different magnitudes (Schweimer and Hauber, 2006; Walton et al., 2003; Walton et al., 2002). Investigations where rats chose between qualitatively different options following ACC lesions have yielded mixed results with reports of both null effects (Schweimer and Hauber, 2005) and decreased effort in the context of choice (Hart et al., 2017).

Here, hM3Dq and hM4Di receptors were expressed under a CaMKIIα promoter, putatively targeting primarily excitatory pyramidal neurons (Nathanson et al., 2009; Wang et al., 2013), in contrast with prior studies using lesions or inactivations that silence all neural activity. ACC likely exerts its effects on choice behavior via projections to downstream targets, the densest of which are to dorsal striatum and mediodorsal thalamus (Vogt and Paxinos, 2014), though ACC also sends sparser efferents to ventral striatum and amygdala (Gabbott et al., 2005).

Though reliable, the magnitude of effect observed here with DREADDs was smaller than what we previously observed following lesions (Cohen’s d=1.39 lesions vs d=0.28 G_i_ vs d=0.29 G_q_) (Hart et al., 2017), and also smaller than effects we have previously reported following pharmacological inactivation (Hart and Izquierdo, 2017) and drug exposure (Hart et al., 2018; Thompson et al., 2017). The smaller effect obtained with DREADDs could be due to different factors. First, DREADD receptors target only cells that recognize the CaMKIIα promoter (putative excitatory projection neurons). Second, although our slice experiments have shown that DREADD receptors modulate neural activity in ACC, these changes in neural activity may have weaker effects on behavior than complete pharmacological inactivation of ACC, or following chronic psychostimulant exposure (Hart et al., 2018). Nevertheless, the DREADDs manipulations were enough to significantly bias behavior away from the preferred, effortful option during choice sessions.

### ACC calcium imaging

ACC interference affected lever pressing only during choice sessions. We used *in vivo* calcium imaging to observe how ACC neurons responded to lever pressing and reward retrieval during such PR, PRC, and CON sessions. A total of 227 neurons from 4 rats were successfully recorded during at least one session of each type and had a significant response to lever pressing or head entry. About two-thirds of these neurons (162/227) were lever-press responsive in at least one of the three session types. Many of these cells (96/162 cells) were also significantly responsive to head-entries in other sessions, indicating some cells maintain encoding properties while others alter response characteristics during different sessions. Calcium trace activity during pre-lever responses was not significantly different in CON compared to PR sessions. However, calcium activity during the pre-lever period was lower during PRC than both PR and CON sessions, suggesting that free chow availability, not satiety from chow, attenuated pre-lever calcium activity.

Another two-thirds of recorded ACC neurons (161/227) were reward-responsive. These sucrose responses were similar in magnitude during CON and PR sessions, indicating that satiety from prior chow consumption had little effect on ACC responses to sucrose. By contrast, sucrose-related calcium activity was significantly lower during PRC sessions than during PR and CON sessions, demonstrating that independently of satiety state, free chow availability attenuated sucrose responses of ACC neurons below the levels seen when sucrose was earned in the absence of free chow.

We next evaluated whether single cell responses reflected the overall population average, but found very few cells that individually responded accordingly in their PETHs (i.e. PR > PRC, CON>PRC, PRC=CON). This implies that the population signal which disambiguates effortful contexts is an emergent property of many cells functioning independently, rather than as a homogenous population. This finding is not surprising given ACC’s role in diverse behaviors and in line with recent single-unit recordings within an effortful task demonstrating robust heterogeneity of response characteristics (Porter et al. 2019). We further looked to see if individual neurons were evenly distributed among conditions types (PR, PRC, and CON). For both LP and HE responsive cells, we found that many cells responded significantly in all 3 session types. Therefore, while individual average responses do not reflect the same population profile, individual cells are preferentially active across similar contexts and serve to disambiguate relative reward value through coordinated population activity. It has been theorized that such mixed selectivity in cortical regions is important in generating high dimensional representations for complex, adaptive behavior (Fusi et al., 2016). This type of heterogeneous population code would be more susceptible to perturbation by either bulk inhibition or excitation, as demonstrated in the results of our DREADDs manipulations.

These findings converge on the idea that activity of ACC ensembles during sucrose consumption encodes the difference between the value of the sucrose versus the value of other available reward options. If no other options are available (as during PR and CON sessions), then the value of other options is zero, and thus nothing is subtracted from the value of the sucrose reward. But if a lower value reward option (such as lab chow during PRC sessions) is available while the rat is working for sucrose, then the nonzero value of the other option may be subtracted from the value of the sucrose reward, reducing the magnitude of ACC responses during sucrose delivery. If this relative value signal for sucrose in ACC is involved in driving motivated effort to work for sucrose, then it would be expected that the rat should exert less effort for sucrose under conditions where the ACC responses to sucrose are smaller. This is exactly what we observed: sucrose responses of ACC neurons were lower during PRC sessions (where lab chow was available as a competing reward option) than during PR or CON sessions (where sucrose was the only reward option).

In summary, neural activity associated with sucrose pellet collection in ACC is strongest when sucrose is the only available option, and weakened by the presence of the counterfactual choice (Mashhoori et al., 2018) (Blanchard and Hayden, 2014), or the value of leaving a patch in pursuit of another option (Hayden et al., 2011). We add here the novel mechanism that ACC modulates this evaluation of options with a subpopulation of stable-coding neurons, which we harnessed the power of calcium imaging to reliably track. Overall, ACC responses to lever pressing and reward-retrieval were lower during PRC sessions; therefore, a plausible explanation of our interference experiments would be that this lower, heterogenous population activity is more susceptible to interference, thus explaining why CNO reduced lever pressing selectively in PRC sessions.

### ACC inhibition versus stimulation

If ACC activity encodes relative value signals that are involved in deriving an animal’s motivation to exert effort, then it is natural to predict that disrupting ACC activity at the neural level would alter effort exertion at the behavioral level. We also expected that bidirectional manipulations of neural activity might yield bidirectional effects upon motivated effort, like manipulations of dopamine (Farrar et al., 2010; Nunes et al., 2013; Randall et al., 2014a; Yohn et al., 2015a; Yohn et al., 2015b). Enhancing dopamine transmission with major psychostimulants (Yohn et al., 2016c), dopamine transporter blockers (Randall et al., 2014b; Yohn et al., 2016a), adenosine A2A receptor antagonists (Randall et al., 2012), or 5-HT2C ligands (Bailey et al., 2018; Bailey et al., 2016) can increase effort output in otherwise untreated rats.

We applied a similar rationale for our G_q_ and G_i_ DREADD experiments, and tested whether ACC chemogenetic inhibition versus stimulation yielded bidirectional effects upon behavioral responding. This was not the case: lever pressing behavior during PRC sessions was similarly attenuated by both G_q_ and G_i_ DREADDs. Indeed, the contributions of ACC and other frontocortical regions to effort may be more complex: pharmacological stimulation of orbitofrontal cortex decreases PR responding (Munster and Hauber, 2017), and GABA antagonism of infralimbic cortex similarly decreases high-effort choice (Piantadosi et al., 2016). Therefore, in frontal cortex there may be an optimal excitatory/inhibitory ratio for computing relative cost-benefit and, consequently, sending appropriate output to downstream targets. The results of ACC stimulation here are consistent with our manipulation introducing noise to otherwise normal neural computations (Mainen and Sejnowski, 1995; Stein et al., 2005), thus impairing behavior in a manner similar to when neural activity is inhibited. Lever pressing rates and task-evoked activity levels in ACC were both lower during PRC sessions than PR or CON sessions. Consequently, disruptions in decision making may occur via changes in signal-to-noise ratio in this region: either by decreases in the signal (i.e. G_i_ DREADDs), or by increases in background noise (i.e. G_q_ DREADDs).

In conclusion, our findings suggest that the role of ACC in effort-based choice may be to discriminate the utility of available choice options by providing a stable population code for the relative value of different reward options. A better understanding of ACC contributions to effort-based choice may yield insight into the mechanisms underlying motivational symptoms in depression (Nunes et al., 2013) and addictions (Robinson et al., 2013).

## Acknowledgements

This work was supported by UCLA’s Division of Life Sciences Recruitment and Retention fund (Izquierdo), R01 DA047870 (Izquierdo), NSF Neuronex 170748 (Blair), the Training program in Translational Neuroscience of Drug Abuse (T32 DA024635, London), and the Training program in Neural Microcircuits (T32 NS058280, Feldman). We thank members of the Izquierdo lab for helpful comments on a previous version of the manuscript. We acknowledge the Staglin Center for Brain and Behavioral Health for additional support related to fluorescence microscopy and UCLA Graduate Division for the Dissertation Year Fellowship (Hart).

## Extended Data

**Figure 3-1.**
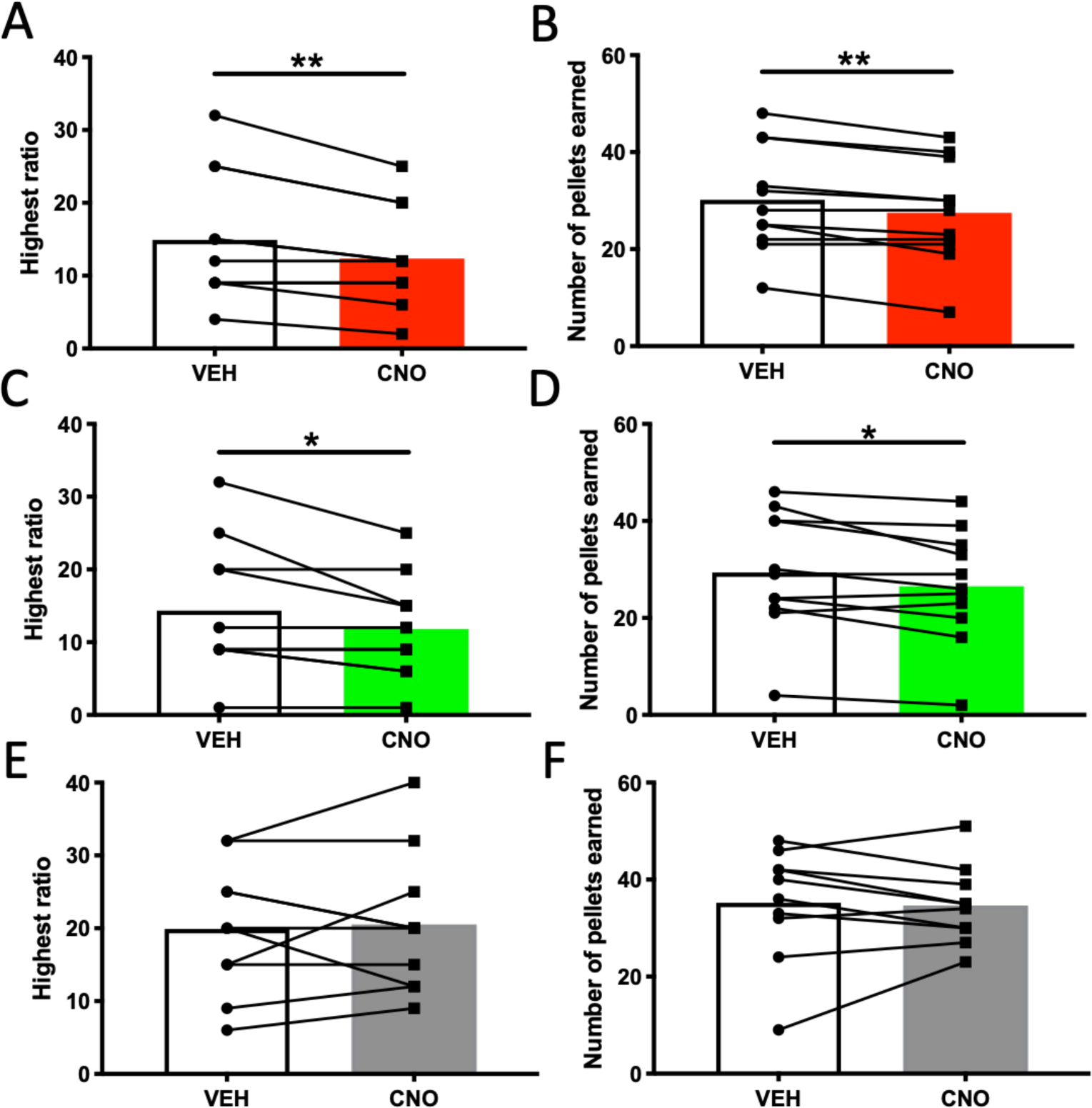
Conventional measures of PR responding during PRC sessions. Highest ratio achieved and number of pellets earned when rats were presented with both the possibility of lever pressing under a PR schedule for sucrose pellets and freely-available chow. Shown is within-subject, counterbalanced choice behavior under vehicle and CNO. **(A)** Highest ratio achieved and **(B)** number of sucrose pellets earned during PRC sessions, when rats were presented with both options in the G_i_ DREADDs condition, in rats receiving CNO compared to within-subject vehicle (VEH). **(C)** Highest ratio achieved and **(D)** number of sucrose pellets earned during PRC sessions, when rats were presented with both options in the G_q_ DREADDs condition, in rats receiving CNO compared to within-subject vehicle (VEH). **(E)** Highest ratio achieved and **(F)** number of sucrose pellets earned during PRC sessions, when rats were presented with both options in the GFP null virus condition, in rats receiving CNO compared to within-subject vehicle (VEH). **p<0.01, ***p<0.001.

**Figure 7-1.**
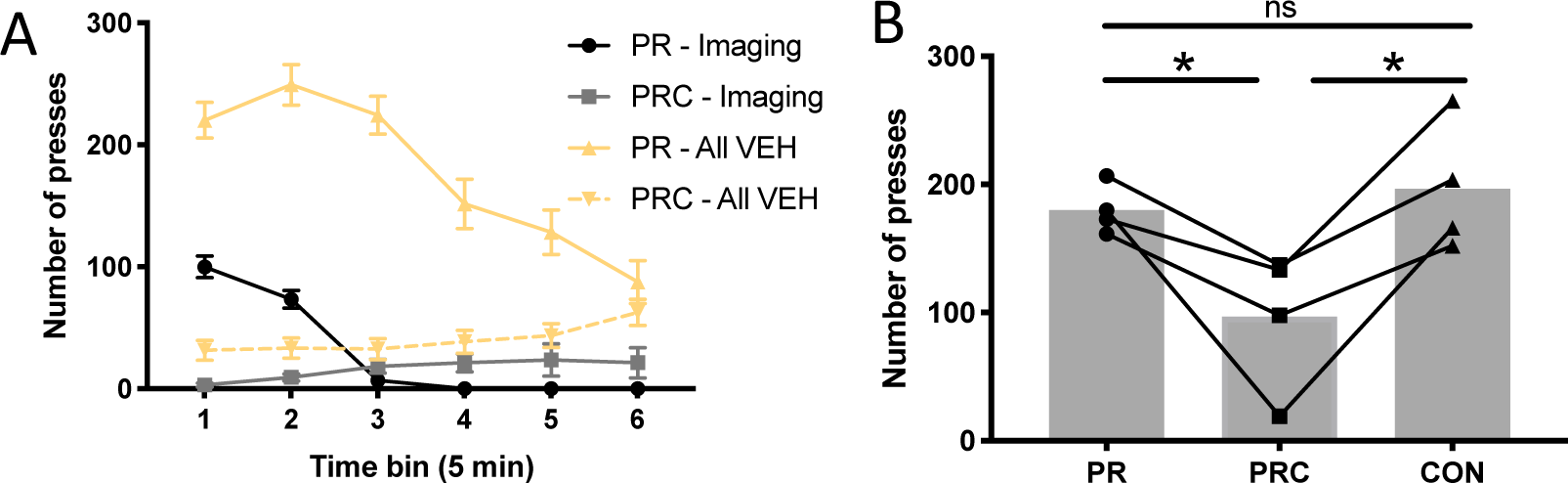
Behavior during imaging sessions. **(A)** Time course of lever pressing behavior in 5 min bins in all imaging sessions, compared to all VEH sessions from the DREADDs experiments. Imaging animals are plotted in grey. DREADDs animals are plotted in gold. **(B)** Average total number of lever presses emitted per session across all imaging sessions.

**Figure 8-1.**
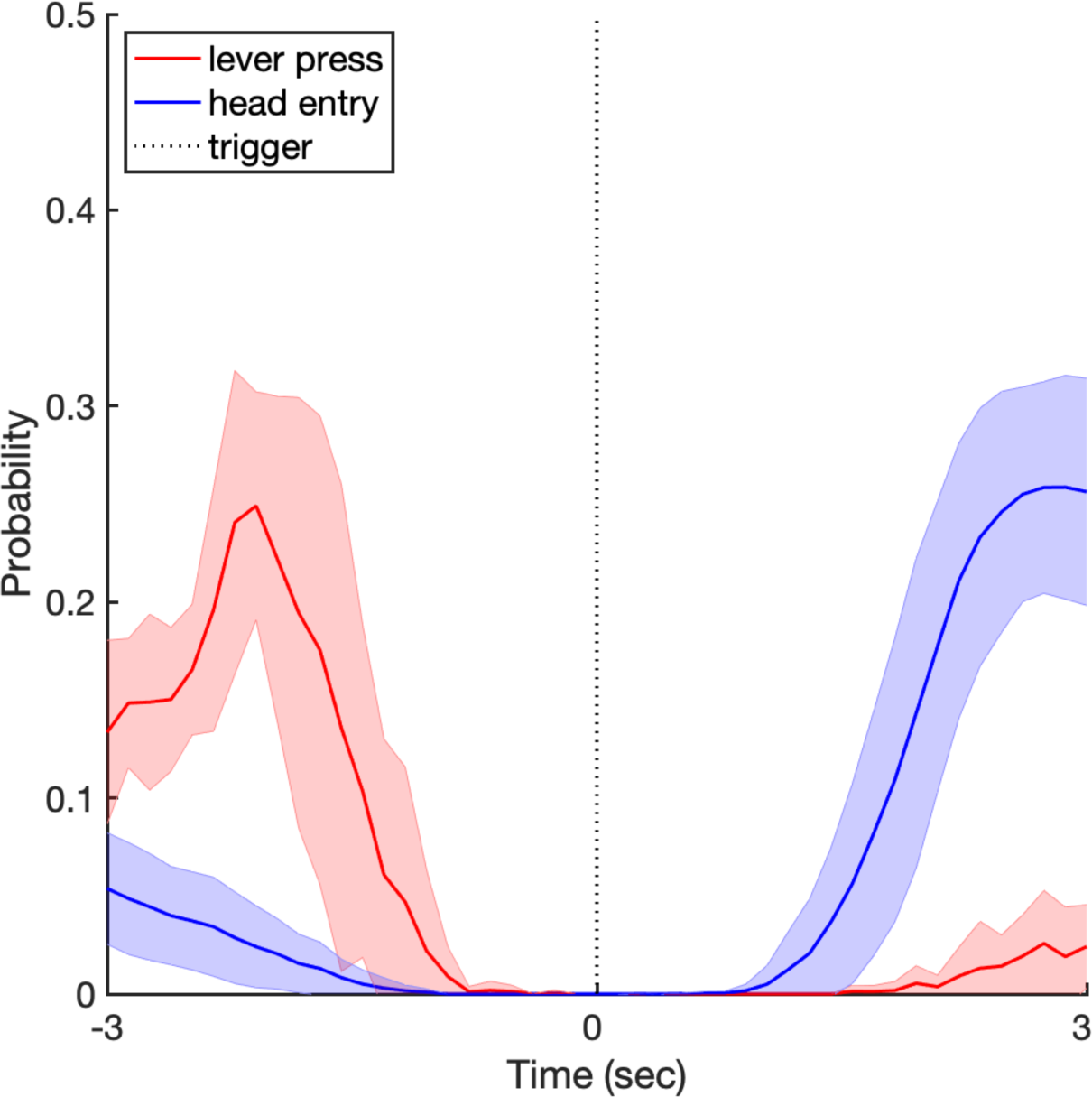
Behavior triggered behavior PETH demonstrating behavioral overlap in post-LP and pre-HE periods. Histogram of HE triggered by LP (red), and LP triggered by HE (blue). The 3 s time window following LP shows substantial contamination by HE events, and the 3 second window prior to HE shows contamination by LP. Contaminated time windows were thus excluded from our analysis. Shaded regions denote 1 ±SEM

